# Synthetic integrin antibodies discovered by yeast display reveal αV subunit pairing preferences with β subunits

**DOI:** 10.1101/2024.01.26.577394

**Authors:** Yuxin Hao, Jiabin Yan, Courtney Fraser, Aiping Jiang, Murali Anuganti, Roushu Zhang, Kenneth Lloyd, Joseph Jardine, Jessica Coppola, Rob Meijers, Jing Li, Timothy A. Springer

**Affiliations:** Program in Cellular and Molecular Medicine, Department of Pediatrics, Boston Children’s Hospital, Boston, MA, USA; Department of Biological Chemistry and Molecular Pharmacology, Harvard Medical School, Boston, MA, USA; Institute for Protein Innovation, Harvard Institutes of Medicine, 4 Blackfan Circle, Room 921, Boston, MA 02115

## Abstract

Eight of the 24 integrin heterodimers bind to the tripeptide Arg-Gly-Asp (RGD) motif in their extracellular ligands, and play essential roles in cell adhesion, migration, and homeostasis. Despite similarity in recognizing the RGD motif and some redundancy, these integrins can selectively recognize RGD-containing ligands including fibronectin, vitronectin, fibrinogen, nephronectin and the prodomain of the transforming growth factors to fulfill specific functions in cellular processes. Subtype-specific antibodies against RGD-binding integrins are desirable for investigating their specific functions. In this study, we discovered 11 antibodies that exhibit high specificity and affinity towards integrins αVβ3, αVβ5, αVβ6, αVβ8, and α5β1 from a synthetic yeast-displayed Fab library. Of these, 6 are function-blocking antibodies containing an R(G/L/T) D motif in their CDR3 sequences. We report antibody binding specificity, kinetics, and binding affinity for purified integrin ectodomains as well as intact integrins on the cell surface. We further employed these antibodies to reveal binding preferences of the αV subunit for its 5 β-subunit partners: β6=β8>β3>β1=β5.

## Introduction

Integrins are critical non-covalent heterodimeric cell surface receptors required for cell adhesion, migration, and signaling. They function as bidirectional signaling molecules by binding to extracellular ligands and intracellular adaptors to the actin cytoskeleton to regulate integrin activation and downstream signaling^1–3^. There are 24 known integrin heterodimer pairs formed by 18 α subunits and 8 β subunits. Eight are RGD-binding integrins that interact with the Arg-Gly-Asp (RGD) motif in extracellular ligands, thereby regulating diverse pathological processes^4–9;10^. αVβ1, αVβ3, αVβ5, and α5β1, expressed on endothelial cells and fibroblasts, bind to fibronectins, exhibiting overlapping functions in cell spreading and migration^11,12^; αVβ6 and αVβ8 promote TGF-β activation subsequent to binding to the RGD-like motifs in prodomain^13^; αIIbβ3 on platelets binds to fibrinogen, playing a critical role in hemostasis^14^; and α8β1 binds to nephronectin in the extracellular matrix and regulates kidney development^15^.

Monoclonal antibodies, peptidomimetics, and small molecule antagonists against RGD-binding integrins have been continuously developed to address the role of integrins in cellular processes^16–18^. However, the similar ligand binding sites among RGD-binding integrin pairs, such as αVβ3 and αVβ5^5^ and αVβ6 and αVβ8^8–10^, pose a significant challenge to development of antibodies that selectively block binding of small, RGD-like ligands. This emphasizes the need to urgently develop molecules specific to RGD-binding integrin subtypes, enabling the discrimination of individual integrins and inhibiting ligand binding at the cellular level. Such advancements are essential to unravel the distinctive biological functions of these integrins and expedite drug development.

Synthetic antibody libraries^19–21^ have distinctive features that we hypothesized could be beneficial in obtaining function-blocking antibodies to integrins. In contrast to traditional species-specific monoclonal antibodies, synthetic libraries can be more effective for selecting antibodies targeting both human and mouse antigens, especially when aiming at highly conserved antigens across different species or conserved sites such as those for ligand binding. Yeast or phage Fab libraries are effective in generating antibodies towards highly conserved proteins, as they do not rely on the self-tolerance mechanisms of the immune system. These libraries typically encode a larger number of unique sequences than the number of B lymphocytes in laboratory animals. In addition, synthetic libraries offer other advantages, such as shorter turnaround times and greater scalability.

Yeast synthetic Fab libraries have the merits of the enhanced protein quality control of eukaryotic cells and suitability for fluorescence-activated cell sorting (FACS) and magnetic-activated cell sorting (MACS) compared to phage libraries^22^. However, the key determinant for successful antibody selection from the yeast display platform is the availability of high-quality antigens. The ectodomains of membrane proteins such as integrins are glycosylated and disulfide-linked, requiring expression in mammalian cells. A non-profit organization, the Institute for Protein Innovation (IPI), has established an antibody platform constructed around yeast display technology, enabling the discovery of antibodies with defined properties. We collaborated to identify antibodies that specifically target RGD-binding integrins, including 6 antibodies containing R(G/T/L) D motifs in their complementarity determining region (CDR)3 with inhibitory functions. Most function-blocking antibodies against integrins do not bind to the ligand binding pocket and block only macromolecular ligand binding due to steric hindrance^4,23,24^; in contrast, antibodies described here are capable of blocking the binding of small molecule, peptidomimetic integrin inhibitors, as well as biological ligands. Several of these antibodies have previously been used to achieve integrin specificity in single molecule studies of integrin force exertion on RGD peptides^12^. To open the way for their usage in integrin biology, and to study how particular assays, conformation dependence, and avidity affect the behavior of these antibodies, we have compared them in multiple assays. As an example of one biological application, we utilized them in investigating the pairing preference of integrin αV subunit with 5 different β subunits and found a consistent preference hierarchy for αV-β pairing on the cell surface.

## Results

### Discovering integrin heterodimer-specific antibodies

We selected for antibodies to RGD-binding integrins αVβ3, αVβ5, αVβ6, αVβ8, and α5β1 using a synthetic yeast-displayed Fab library containing ∼10^10^ unique Fab sequences^12^. We enriched yeast clones displaying integrin-specific Fabs through magnetic-activated and fluorescent-activated cell sorting. Selection steps included positive selection with target integrin ectodomains, negative selection with poly-specificity reagent (PSR) and untargeted integrins, and human/mouse cross-reactivity. After next-generation sequencing, the most frequent 13 sequences for each integrin target were expressed as human IgG1 for characterization. Initial screening assessed specificity towards intact human or mouse integrins expressed on the cell surface of WT K562 or K562 stable transfectants or Expi293F αV^-^/α5^-^ cell transient transfectants. Each antibody is named according to the integrin with which they were selected followed by a number. Immunofluorescent staining at 50 nM antibody concentration identified 11 antibodies selective for the target integrin (Fig. 1 A, B). Six antibodies contained an R(G/T/L) D motif in their heavy chain CDR3 (Table 1). We also evaluated the cross-reactivity of these antibodies on mouse integrins and found that 10 out of 11 antibodies could bind to the target mouse integrin (Fig. 1C-G); however, specificity toward mouse integrins was lower than for human integrins. This may relate to using the mouse antigen as the last step in our selection process and the lack of counter-selection against other mouse integrins. IPI-αVβ6.4, which contains an RTD motif, crossreacts between αVβ6 and αVβ8 in both human and mouse (Fig. 1A and Fig. 1E). This is interesting, as integrins αVβ6 and αVβ8 share specificity for TGF-β1 and β3 prodomain-growth factor complexes (proTGF-β). In summary, we obtained 11 antibodies that can specifically target integrins αVβ3, αVβ5, αVβ6, αVβ8, and α5β1.

**Figure 1.**
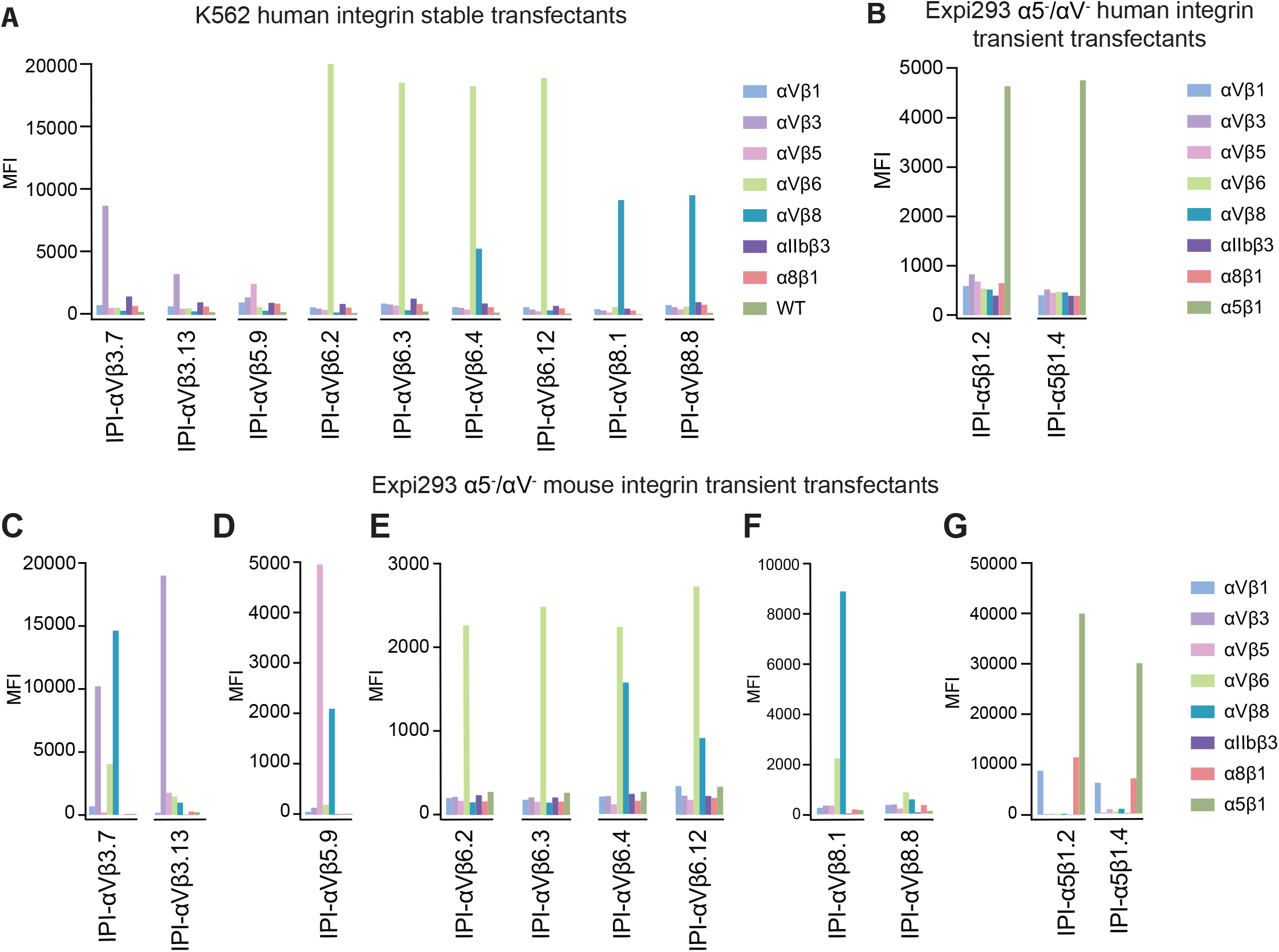
Integrin specificity of antibodies on all RGD-binding human and mouse integrin transfectants by indirect flow cytometry. (A) K562 stable human integrin transfectants in Ca^2+^/Mg^2+^. (B) Expi293 α5^-^/αV^-^ cell transient human integrin transfectants in Ca^2+^/Mg^2+^. (C-G) Expi293 α5^-^/αV^-^ cell transient mouse integrin transfectants in Ca^2+^/Mn^2+^. Immunostaining was performed with 50 nM IPI integrin antibody followed by washing and detection with APC-conjugated goat anti-human secondary antibodies and flow cytometry. MFI: mean fluorescence intensity.

**Table 1.**
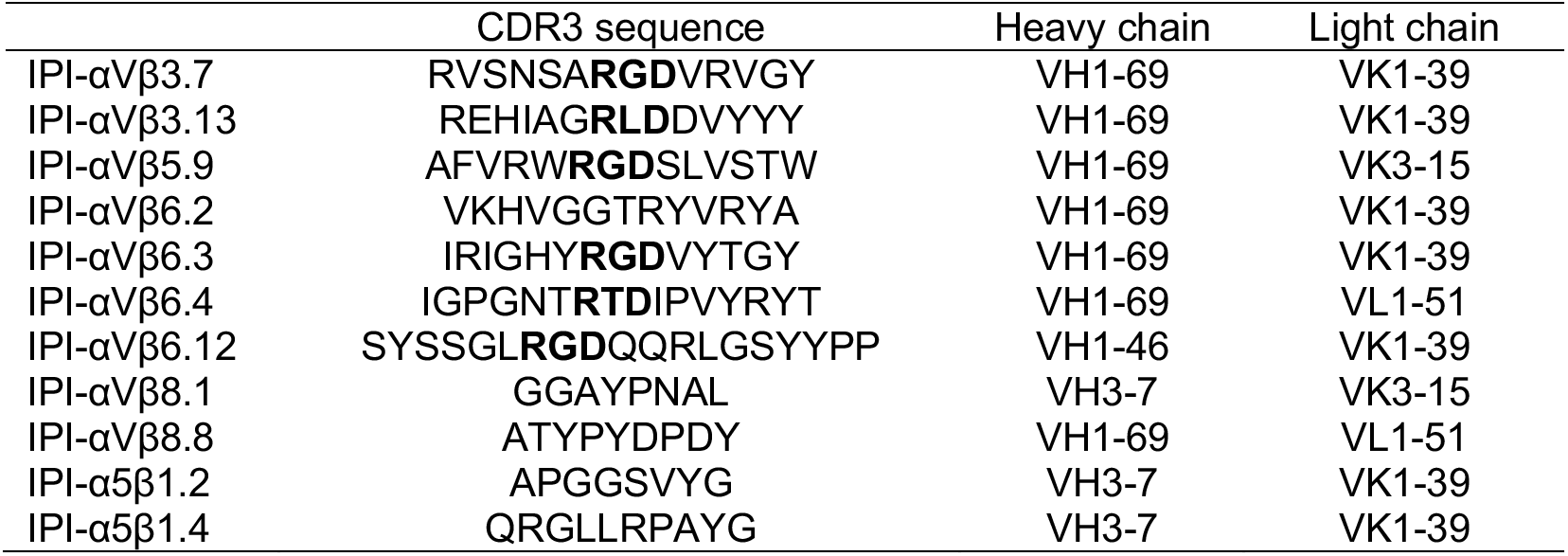
IPI integrin antibody sequence.

We next titrated the antibodies in immunofluorescent staining of K562 stable transfectants or WT K562, which expresses α5β1, using a secondary fluorescent anti-IgG (Fig. 2). For an antibody specific for αVβ1, we used sequence 5 from a Biogen patent^25^, which we designate Biogen-αVβ1.5. The EC50 values ranged from 0.2 to 6 nM.

**Figure 2.**
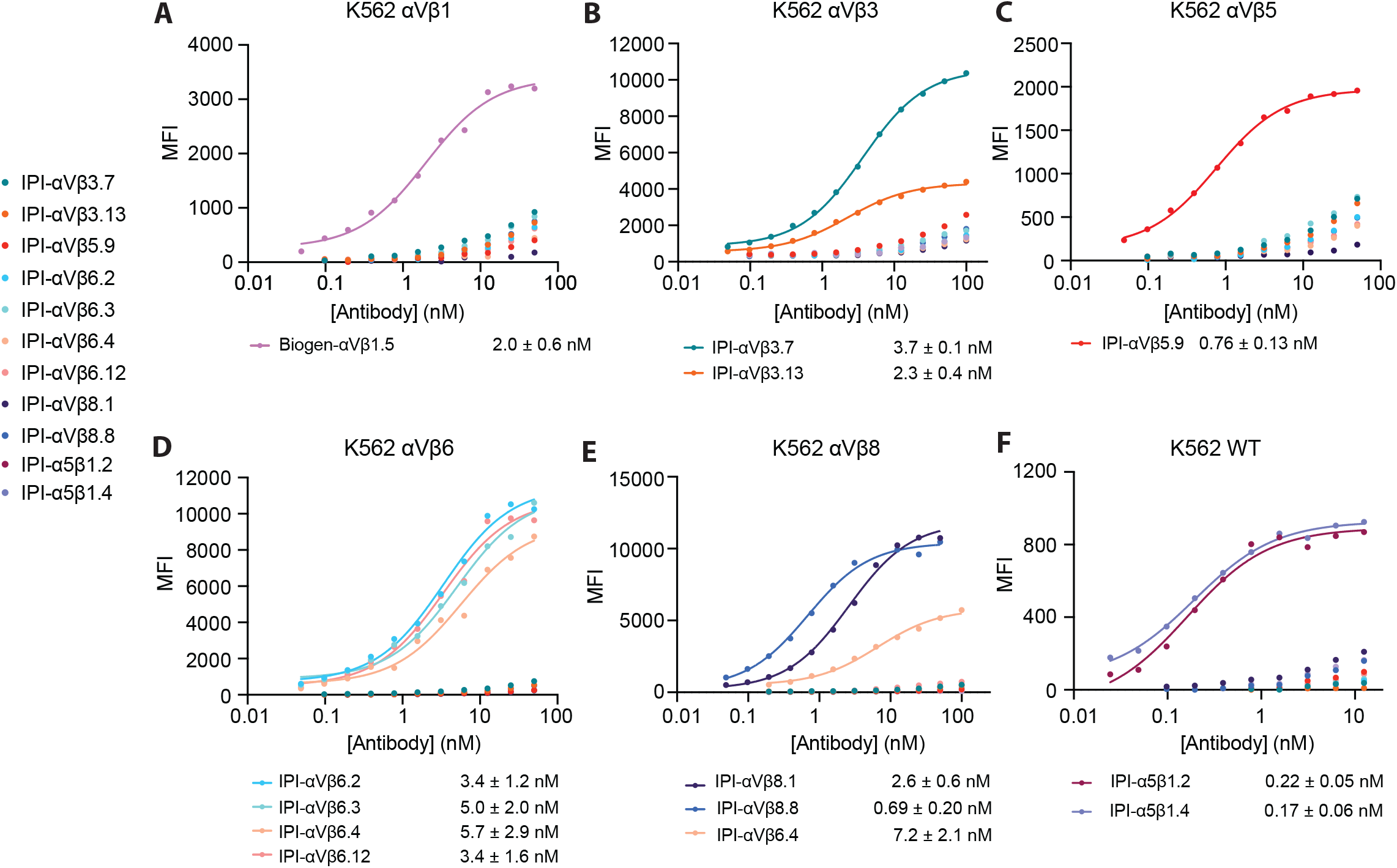
Titration of antibodies on human RGD-binding integrin K562 stable transfectants by indirect flow cytometry. All antibodies were titrated against each transfectant in Ca^2+^/Mg^2+^ and immunostaining was as in Figure 1. The mean fluorescent intensity (MFI) at each antibody concentration after subtraction of isotype control at the same concentration was fitted to a three-parameter dose-response curve for EC50, background MFI, and maximum MFI; curves are only shown for antibodies with meaningful staining. The errors for the EC50 values are the standard errors from the non-linear least square fits.

### Binding kinetics and affinity measurement with surface plasmon resonance (SPR)

We measured the binding of immobilized antibodies to the purified soluble ectodomains of all 8 RGD-binding integrins by SPR (Fig. 3 and Fig. S1-4). All 11 antibodies demonstrated high affinity for their target integrin subtypes, with affinities ranging from sub-nanomolar to two-digit nanomolar (Table S1). The dissociation rate constant (k_off_) values were in the range of 1×10^-4^ to 1×10^-3^s^-1^ with an average of 5.2×10^-4^ s^-1^.

**Figure 3.**
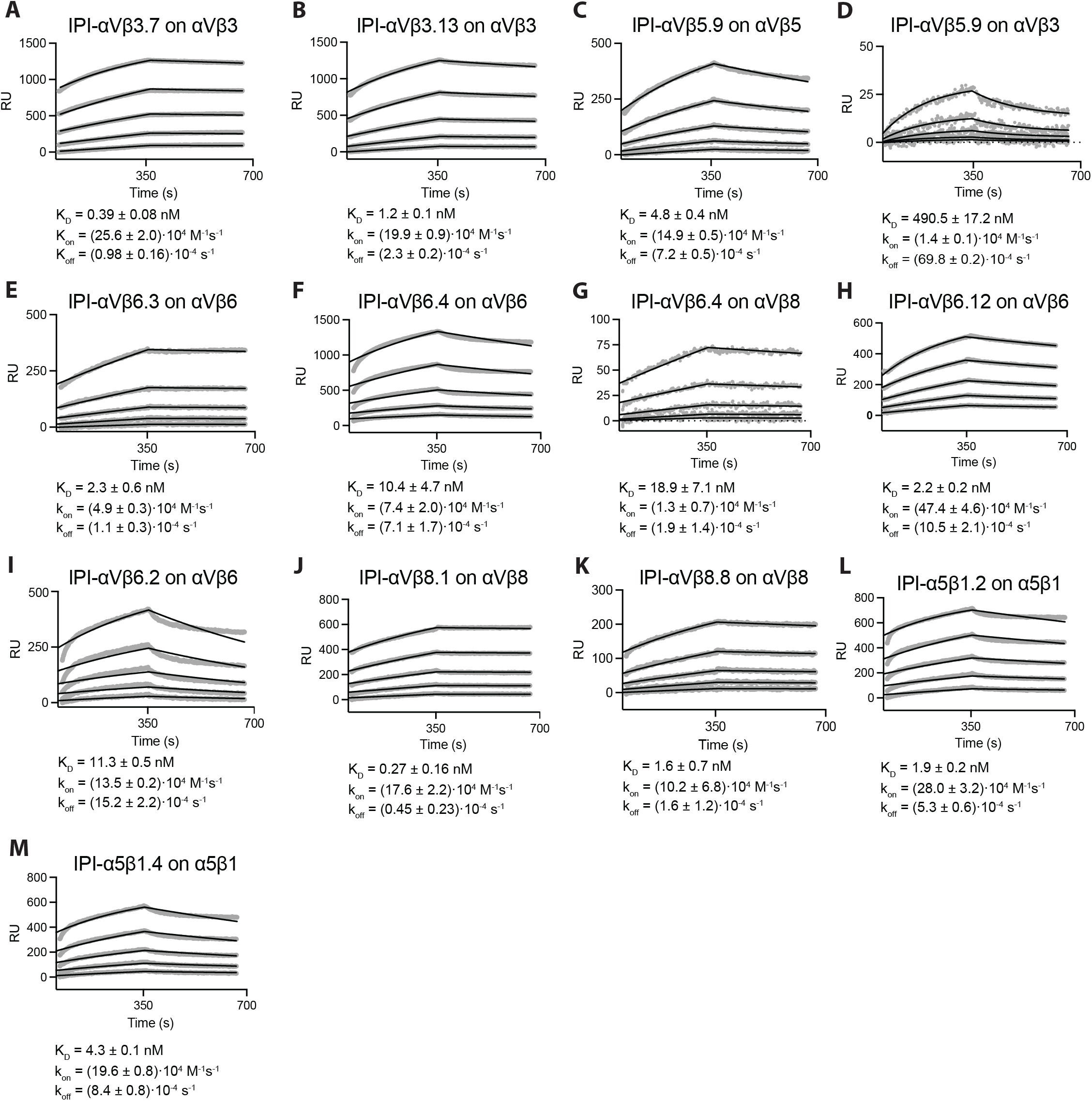
Surface plasmon resonance (SPR) binding kinetics with soluble integrin ectodomains. (A-M). Antibodies were captured on the surface with anti-Fc. Integrins in 10 mM HEPES pH 7.5, 150 mM NaCl, 1 mM MgCl_2_, 1 mM CaCl_2_, 0.05% Tween 20, and 0.5 mg/mL BSA were used at 0.78, 1.56, 3.12, 6.25, and 12.50 nM. SPR sensorgrams (thick gray lines) at each ectodomain concentration were globally fitted with 1 vs 1 Langmuir binding model for the on- and off-rates, k_on_ and k_off_. K_D_ values were calculated as k_off_ /k_on_. Values are reported as means with standard deviations from three independent regions of interest (ROIs).

Most antibodies, including the ones with RGD-like motifs, displayed remarkable selectivity toward the target integrin. Antibody IPI-αVβ6.4, which cross-reacts with mouse and human αVβ8, bound to αVβ8 with ∼2-fold lower affinity than αVβ6 (Fig. 3F, G). Other antibodies with RGD-like motifs cross-reacted with non-cognate integrins with >100-fold lower affinity (Fig. 3D, Fig. S1C, 2A, 2C, and Table S1). Among the five non-RGD-containing antibodies, significant crossreactivity was found only for IPI-αVβ6.2, which bound to αVβ8 with 15-fold lower affinity than αVβ6 (Fig. S3A, Table S1).

### Competitive binding assays with RGD-mimetic antibodies using soluble integrin ectodomains

Solid phase assays, such as SPR, offer advantages but suffer from potential artifacts not present in solution phase assays. Antibody competition with FITC-labeled peptidomimetic ligands in fluorescence polarization (FP) is a solution phase assay, and also allowed us to test the hypothesis that antibodies with RGD-mimetic sequences in their heavy chain CDR3 bound to integrin ligand-binding sites. We measured concentration-dependence of competition by antibodies of binding of fixed concentrations of FITC-labeled, disulfide-cyclized ACRGDGWCG peptide (FITC-cyclic-ACRGDGWCG) or FITC-labeled GRGDLGRLKK peptide (FITC-proTGFβ3 peptide) to a fixed concentration of integrin ectodomain.

All six RGD-mimetic antibodies successfully competed with the FITC-cyclic-ACRGDGWCG or FITC-proTGFβ3 peptide ligands, demonstrating competition at the ligand-binding site (Fig. 4, Fig. S5). Affinities for the target integrin ectodomains ranged from 0.7 to 11.3 nM. Competition by all antibodies with both peptide ligands revealed cross-reactivity among RGD-binding integrins for some RGD-mimetic antibodies, but with affinities hundreds to thousands times lower than to the target integrins. For example, IPI-αVβ5.9 had 700-fold lower affinity for αVβ3 than αVβ5 (Fig. 4B and C). IPI-αVβ6.12 bound to αVβ3 and αVβ8, with affinities 1000-fold and 300-fold lower, respectively, than to its target αVβ6 (Fig. 4B, E, and F). IPI-αVβ3.7 bound to αVβ8 with an affinity 3000-fold lower than to its target, αVβ3 (Fig. 4B and F).

**Figure 4.**
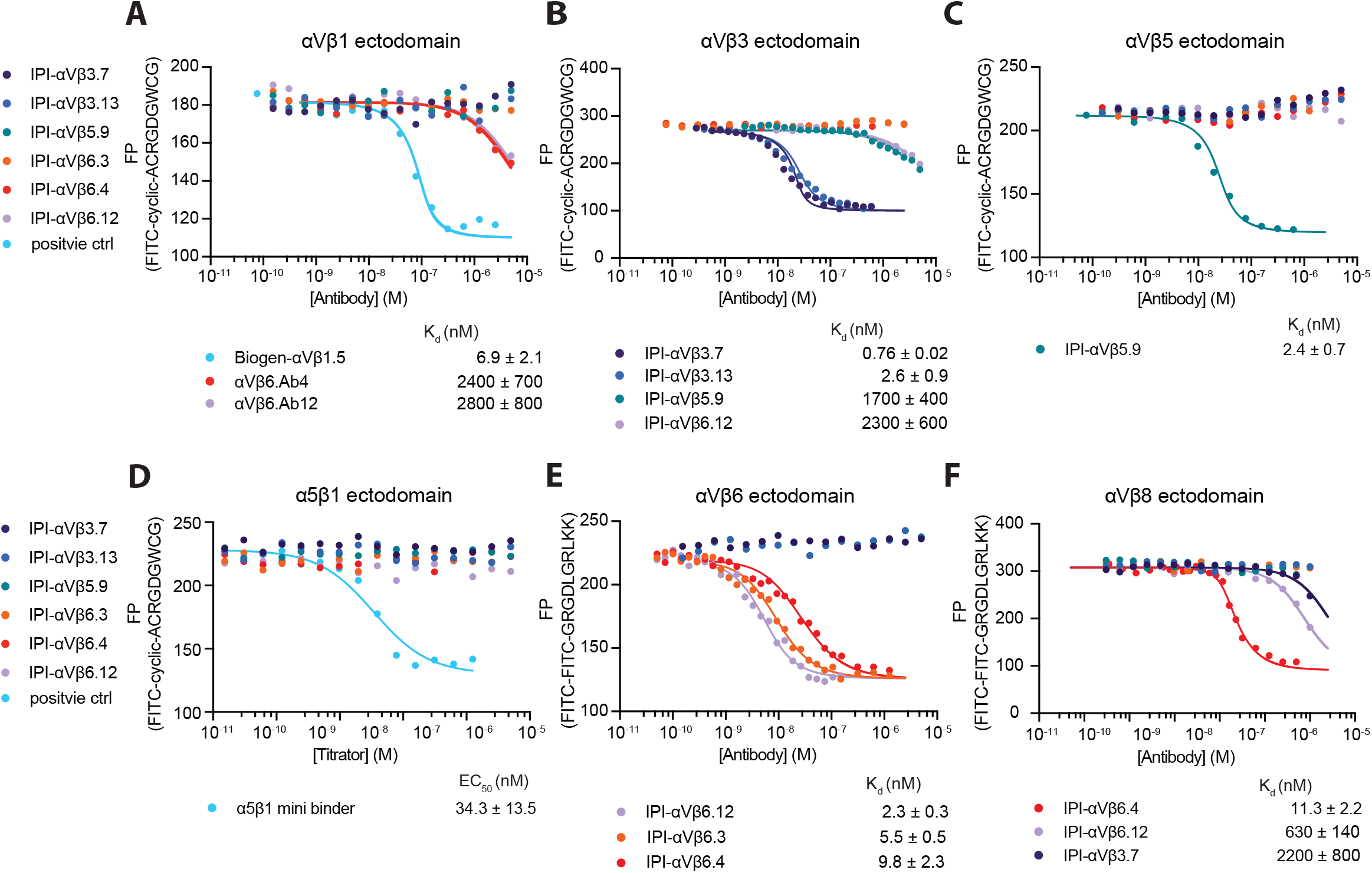
Binding affinities calculated from competition by RGD-mimetic antibodies of ectodomain binding to fluorescent RGD peptides using fluorescence polarization. (A-D) Competition of 10 nM FITC-cyclic-ACRGDGWCG binding to 200nM αVβ1, 50nM αVβ3, 50nM αVβ5 or 100nM α5β1. (E-F) Competition of 10nM FITC-proTGFβ3 peptide binding to 10nM αVβ6 or 200nM αVβ8. Competitive antibody binding curves were globally fitted^26^ with the maximum FP value in the absence of antibody and the minimum FP value as global fitting parameters, and K_D_ value for each antibody as individual fitting parameter (Methods). A reliable fit could not be obtained for the α5β1 minibinder and its EC_50_ value was calculated by fitting the curve with a three-parameter dose-response curve. Means and standard errors are from nonlinear least square fits.

### The effect of avidity on apparent affinity of bivalent RGD-mimetic antibodies for cell surface integrins

Typical immunofluorescence flow cytometry, whether done with a primary or secondary fluorescent antibody, is done with washing and is thus not an equilibrium measurement of affinity (e.g., Fig. 2). True equilibrium measurements of binding of fluorescent ligands can be done by flow cytometry without washing but are challenging at concentrations above 100 nM because of the large excess of free ligand^26^. In this section, we worked around this limitation by measuring cell-bound fluorescence of a fixed concentration of a conformational reporter or RGD mimetic, while titrating in IgG or Fab of RGD-mimetic antibodies.

We first measured the equilibrium affinities and specificities of the RGD-mimetic IgG for cell surface integrins (Fig. 5 and Fig. S6). Binding to β1 integrins was measured by enhancement of binding of Alexa647 labeled 9EG7 Fab, which is specific for the extended states of β1 integrins. None of the six RGD-mimetic antibodies showed detectable binding to intact αVβ1, α8β1, and α5β1 up to 2 μM, while Biogen-αVβ1.5 and cRGD peptide served as positive controls (Fig. 5A-C). Affinities for the other RGD-binding integrins were determined by competing with fluorescently labeled cRGDfK peptide for integrin αVβ3 and αVβ5, proTGF-β3 peptide for integrins αVβ6 and αVβ8, and echistatin for integrin αIIbβ3 (Fig. 5D-H). All six RGD-mimetic antibodies exhibited high affinities ranging from 0.5 to 1.2 nM to the target cell surface integrin. Selectivity was also very high, with no antibodies showing cross-reactivity except for IPI-αVβ5.9, which bound to αVβ3 and αVβ8 with 1.2 μM and 5.2 μM affinity, respectively (Fig. 5D and G).

**Figure 5.**
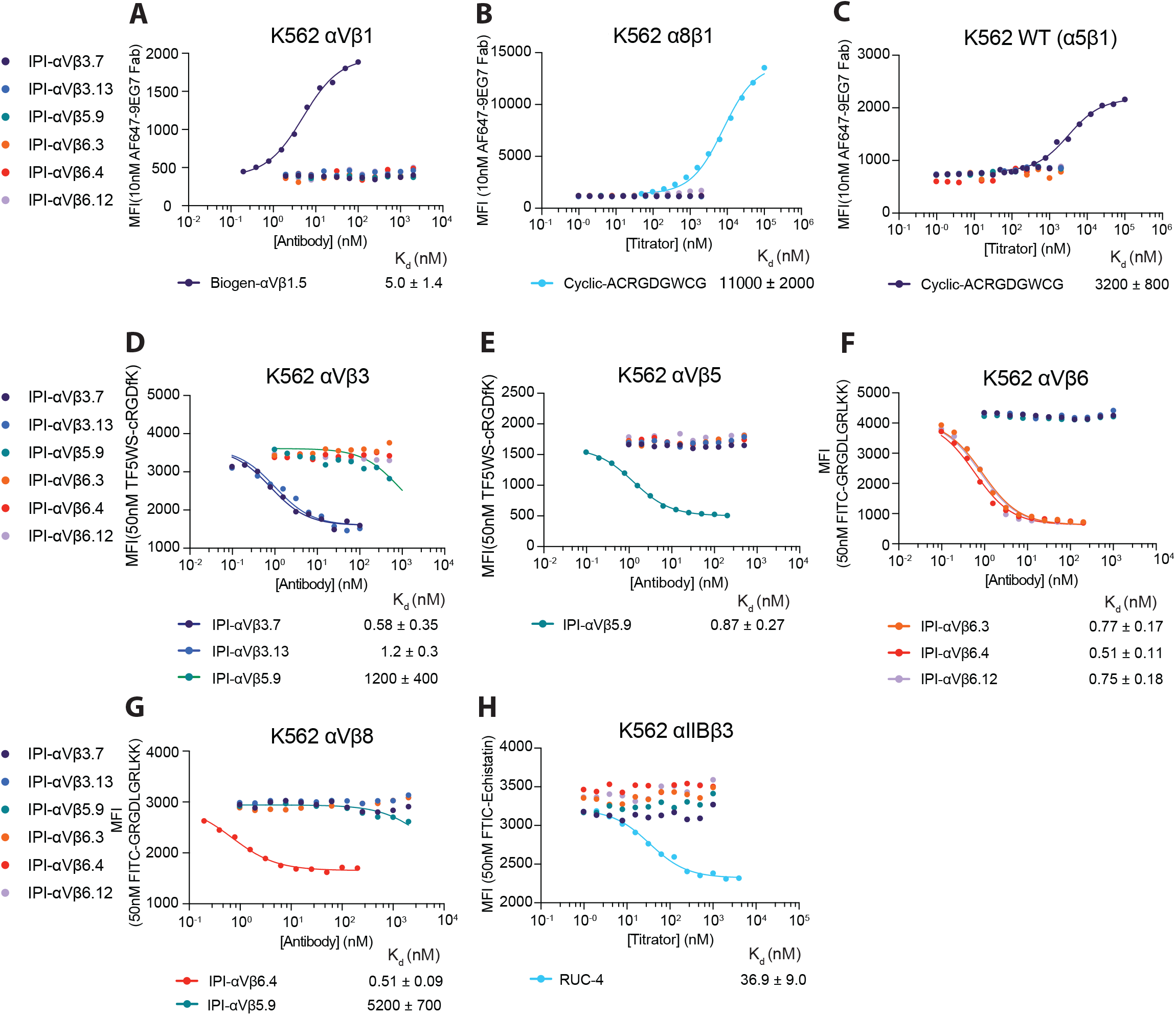
Binding affinities of RGD-mimetic antibodies for cell surface RGD-binding integrins by flow cytometry without washing. (A-C) Affinities on K562 stable transfectants or WT K562 cells were measured by enhancement of binding of 10nM AF647-9EG7 Fab. Cyclic-ACRGDGWCG and Biogen-αVβ1.5 were included as positive contols. Affinities and standard errors are from nonlinear least square fits of MFI values to a three-parameter dose-response curve. (D-H) Affinities on K562 stable transfectants were measured by competing fluorescently labeled RGD-mimetics. Affinities and standard errors are from nonlinear least square fits of MFI values to a three-parameter dose-response curve fitted individually (αVβ5 and αIIbβ3) or fitted globally (αVβ3, αVβ6 and αVβ8) with the minimum MFI and the maximum MFI as shared fitting parameters and EC50 for each titrator as individual fitting parameters. The K_D_ value of each titrator was calculated from the EC50 value as K_D_ = EC50 / (1 + C_L_/K_D,L_), where C_L_ is the concentration of the fluorescent peptidomimetic and K_D,L_ is the binding affinity of the fluorescent peptidomimetic to the respective integrin ectodomain as referenced in Methods. The errors for the affinities are the difference from the mean from duplicate experiments.

We next directly compared the affinities of IgG and Fab (Fig. 6). For all six RGD-mimetic antibodies, IgG bound with higher affinity than Fab. IgG affinity was enhanced from a range of 7.5-fold for IPI-αVβ3.7 (Fig. 6A) to 60 to 70-fold for IPI-αVβ5.9 (Fig. 6B) and IPI-αVβ6.4 (Fig. 6C). Notably, IPI-αVβ6.Ab4 cross-reacts with αVβ8, with which it showed a lesser 27-fold enhancement (Fig. 6D). These results underscore the significant role of avidity effects in binding interactions between these antibodies and cell surface integrins.

**Figure. 6.**
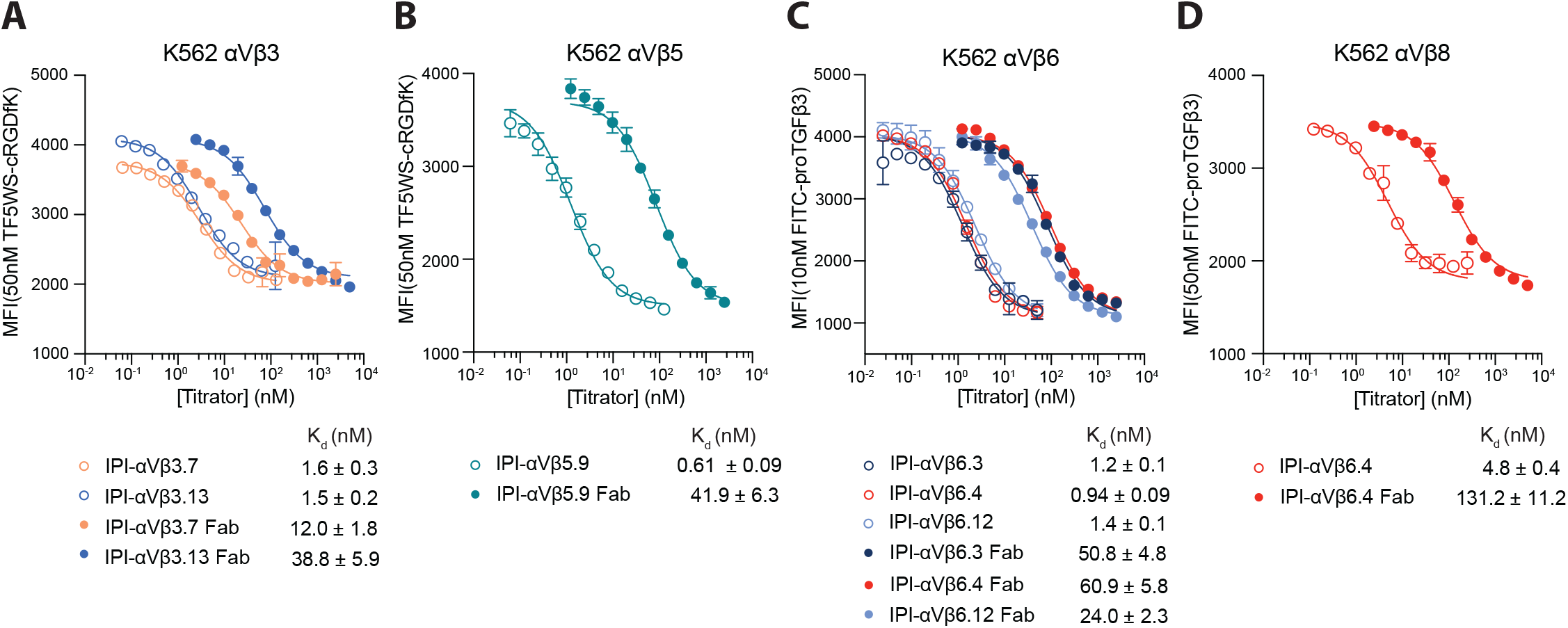
Affinities of RGD-mimetic antibodies and their Fab fragments for cell surface integrins on K562 stable transfectants. Experimental setup and data fitting were as described in Fig. 5.

### Inhibition of integrin-mediated cell adhesion

Many antibodies to integrins inhibit binding to biological ligands by binding to a site adjacent to but not in the RGD-binding pocket. We tested αVβ1, αVβ3, αVβ5 and α5β1-dependent cell adhesion to a fibronectin fragment (Fn3 domains 7-12) and αVβ6 or αVβ8-dependent cell adhesion to proTGF-β1 GARP complexes (Fig. 7). All 6 RGD-mimetic antibodies specifically inhibited integrin-mediated cell adhesion. Additionally, despite lacking a R(G/T/L) D motif, IPI-αVβ6.2 inhibited adhesion to proTGF-β1 GARP complexes (Fig. 7E).

**Figure 7.**
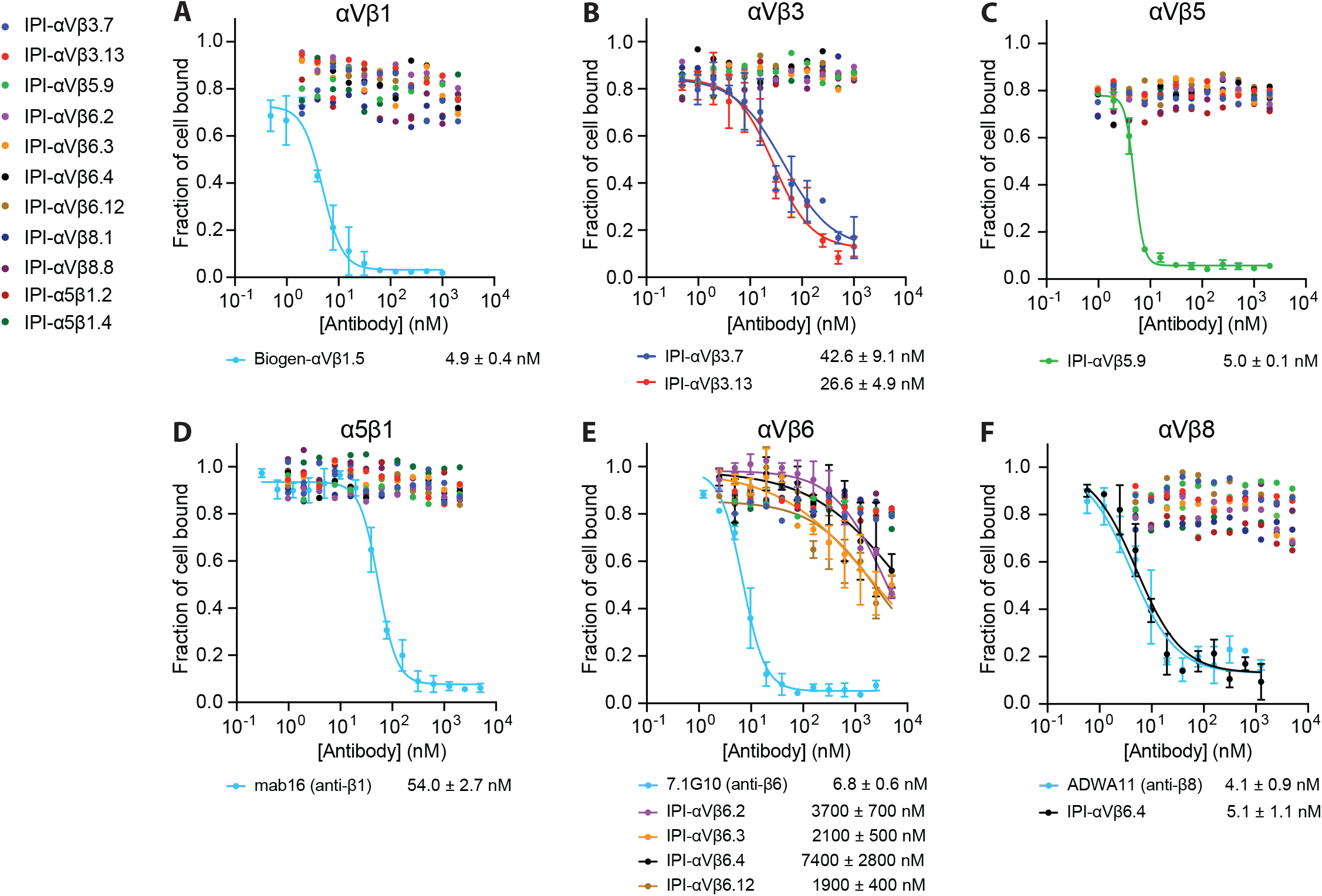
Inhibition of cell adhesion to ligands on substrates. Expi293 αV^-^/α5^-^ KO cells transiently transfected with the indicated integrins were mixed with IPI anti-integrin antibodies and assayed for adhesion to ELISA plates coated with 30 nM fibronectin fragment (Fn3 7-12) (A-D) or with 10 nM GARP ectodomain/proTGFβ1 (E-F). After 1 hr at 37°C, the fluorescent intensity of mCherry, which was co-expressed with the transfected β-subunit using a self-cleaving P2A peptide, was recorded before and after washing away nonadherent cells. The fraction of cells bound at each antibody concentration was fitted individually or globally (if more than one antibody was fitted) to a four-parameter dose-response curve, with global fit to shared bottom and top and individual fit to IC50 and Hill slope. Values are means and s.e. from triplicate measurements.

Most IPI antibodies inhibited adhesion of Expi293 αV^-^/α5^-^ KO transfectants with IC50 values within ∼10-fold of their affinities for cell surface integrins (Fig. 7B, C, and F). However, all four IPI antibodies to αVβ6 inhibited adhesion with far less potency, with IC50 values reduced ∼1,000-fold relative to affinity, while the 7.1G10 antibody^27^ was far more potent (Fig. 7E). In contrast, IPI-αVβ6.4, which cross-reacts with αVβ6 and αVβ8, inhibited αVβ8-dependent adhesion with 1,000-fold more potency than αVβ6-dependent adhesion, and was equipotent to ADWA11 antibody (Fig. 7E and F). The reason for these differences is unclear.

### Pairing preference of αV for the 5 β subunits

Having characterized a set of integrin subtype-specific antibodies, we employed them to investigate whether the αV subunit prefers to associate during biosynthesis with certain of its 5 different β subunit pairing partners over others. To quantify expression we used flow cytometry with fluorescently-labeled integrin heterodimer-specific antibodies. To correct for variation in binding and dissociation kinetics among the antibodies, we multiplied the mean fluorescence intensity (MFI) of each antibody by the ratio of the MFI of αV subunit specific antibody, 17E6, and the heterodimer-specific antibody (Fig. S7).

In preliminary experiments, we determined the optimal αV-β subunit plasmid transfection ratio for each αV heterodimer using Expi293 αV^-^/α5^-^ KO cells to minimize endogenous integrin expression (Methods). The highest expression of αVβ3, αVβ6, and αVβ8 was achieved with equal amounts of αV and β-subunit plasmids (Fig. S7B, D, and E) and of αVβ1 and αVβ5 with αV: β-subunit plasmid ratios of 1:3 (Fig. S7A and C).

To determine pairing preferences, we then used a fixed amount of αV plasmid and varying ratios of β-subunit plasmids (Fig. 8 and Methods). β1 and β5 were outcompeted by all other β-subunits and equally competed with one another (ratio of 0.97); therefore, αVβ1 and αVβ5 are the least favored heterodimers (Fig. 8A-D). β3 outcompeted β1 and β5 (Fig. 8A and E) but in turn was outcompeted by β6 and β8 (Fig. 8F and G). Finally, β6 and β8 competed equally with one another (Fig. 8J). The “pecking order” was therefore αVβ6= αVβ8> αVβ3> αVβ1= αVβ5.

**Figure 8.**
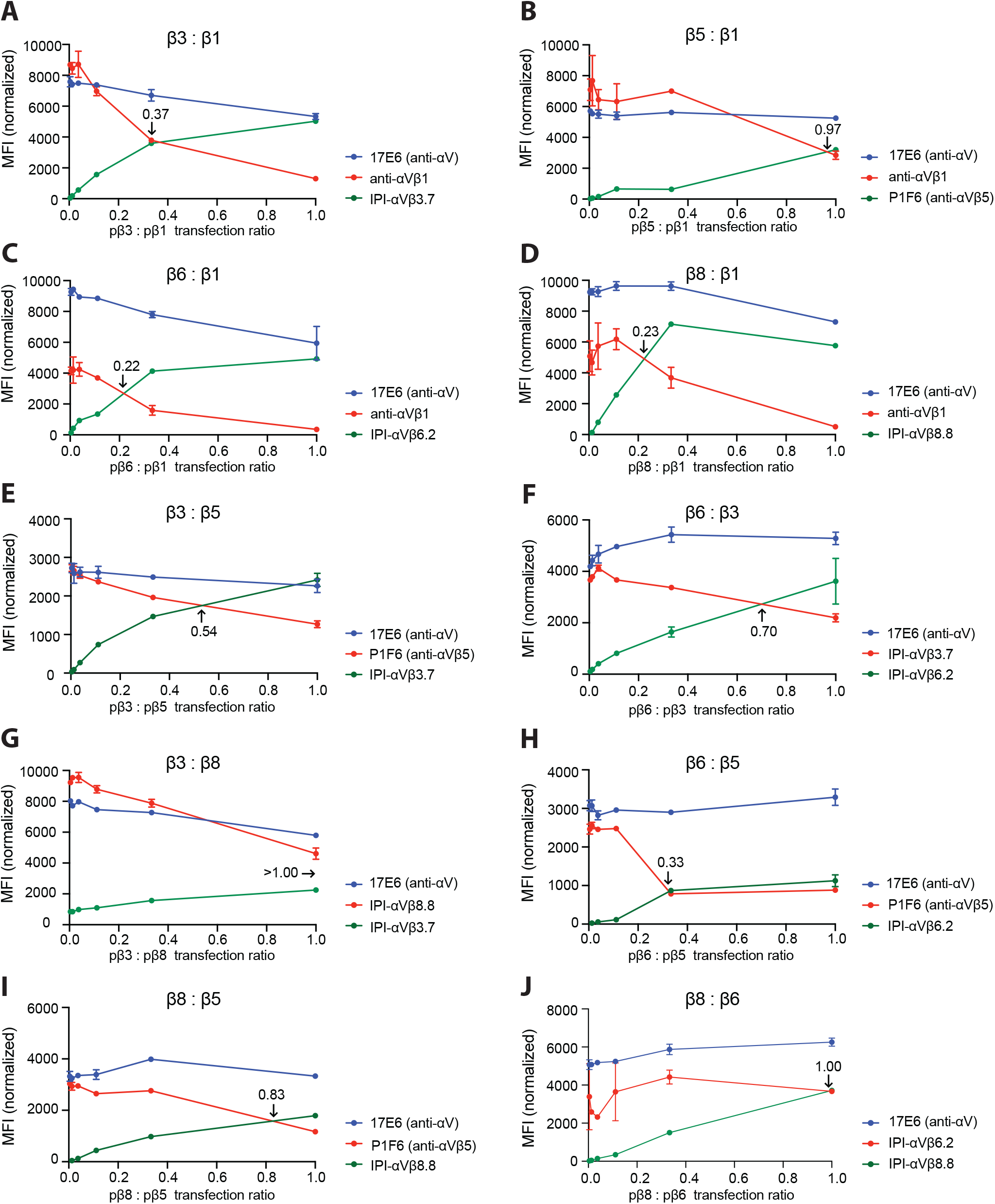
Competition between integrin β-subunits for the αV-subunit. A-J. MFI of directly fluorophore-labeled integrin antibodies measured by flow cytometry. In each competitive titration, the concentration of the αV-subunit plasmid (pαV) and one β-subunit plasmid remained constant at 0.6 µg (red line) while the other β-subunit plasmid (green line) was titrated until reaching 0.6 µg. The αV-subunit plasmid was 0.2 µg in A-E and H-I and 0.6 µg in F-G and J. In all reactions, empty vector plasmid was added to make the total plasmid concentration 1.8 µg. The ratio of the two β subunit plasmids at the cross point is indicated in each panel. The MFI of each β-subunit antibody was normalized relative to the MFI of the 17E6 αV antibody (Fig. S7).

### Integrin αVβ1 heterodimer formation on other cell lines

We extended comparisons among αV integrins to cell lines that express αVβ6 and αVβ8. Glioblastoma cell line LN229 expresses high levels of αV, β1, α5β1, and αVβ3, moderate levels of αVβ5 and αVβ8, no αVβ6, and no αVβ1 (Fig. 9A-C). Colorectal adenocarcinoma cell line HT29 expresses high levels of αV and β1, high levels of αVβ6, moderate levels of αVβ5 and αVβ8, and no αVβ1, αVβ3, or α5β1 (Fig. 9D-F).

**Figure 9.**
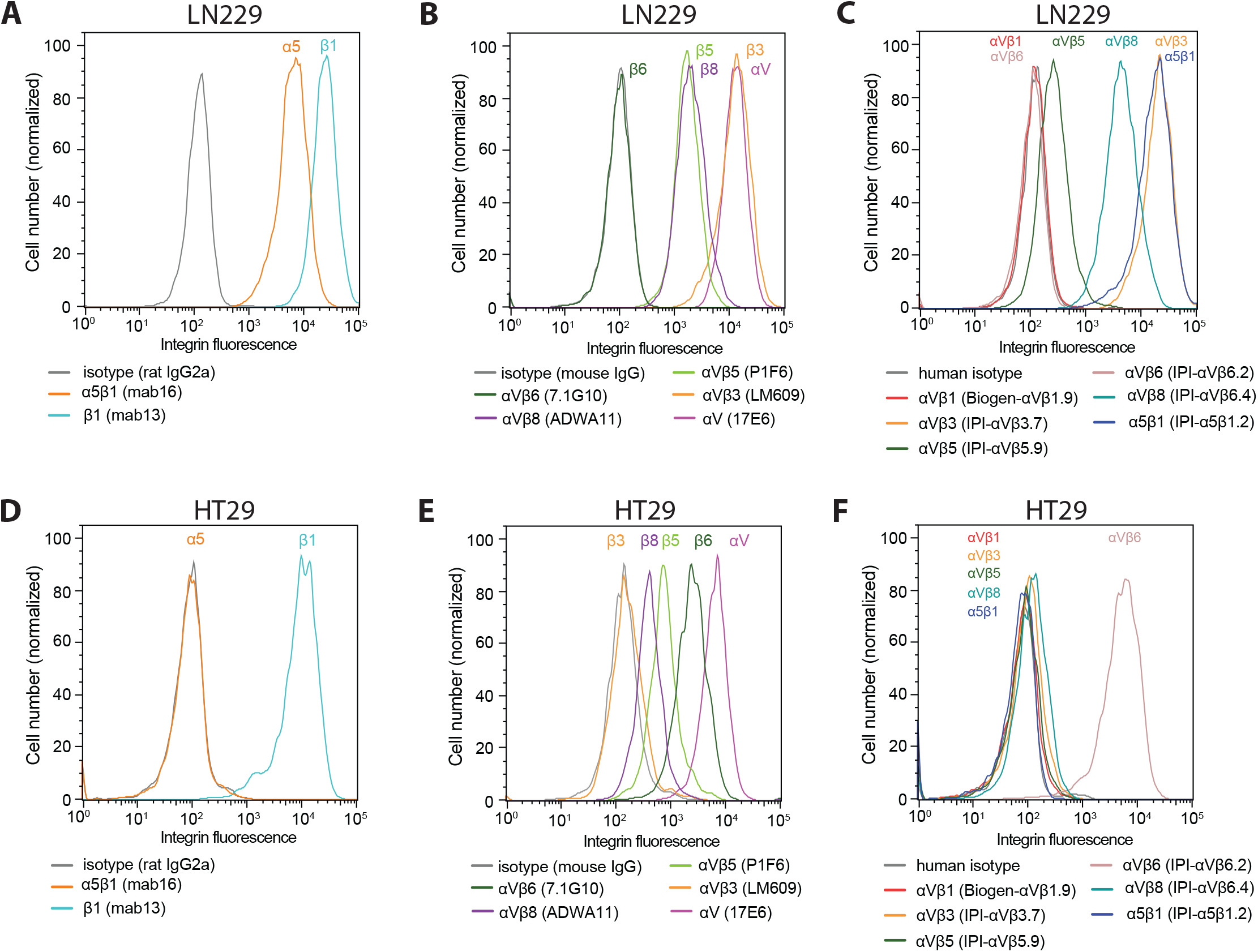
Immunostaining of cell surface integrins on LN229 cells (A-C) and HT29 cells (D-F). Cells were stained with 50 nM of the indicated anti-integrin antibodies or isotype control antibodies in HBSS buffer containing 1 mM Ca^2+^ and 1 mM Mg^2+^ except for IPI-αVβ5.9 which used 1 mM Mn^2+^ and 0.2 mM Ca^2+^. After washing, integrin antibodies were detected using APC-conjugated goat anti-human secondary antibodies, Alexa Fluor 647 goat anti-rat IgG, or Alexa Fluor 647 goat anti-mouse F(ab’)2, and flow cytometry.

## Discussion

We have identified and characterized a suite of antibodies to human integrins, some of which also crossreact with mouse integrins, validated their use in competition with RGD mimetic ligands and in cell adhesion assays (Table 2), and demonstrated their utility in defining the β-subunit preference of the αV integrin subunit. Our data provides guidance for the future application of these antibodies.

**Table 2.**
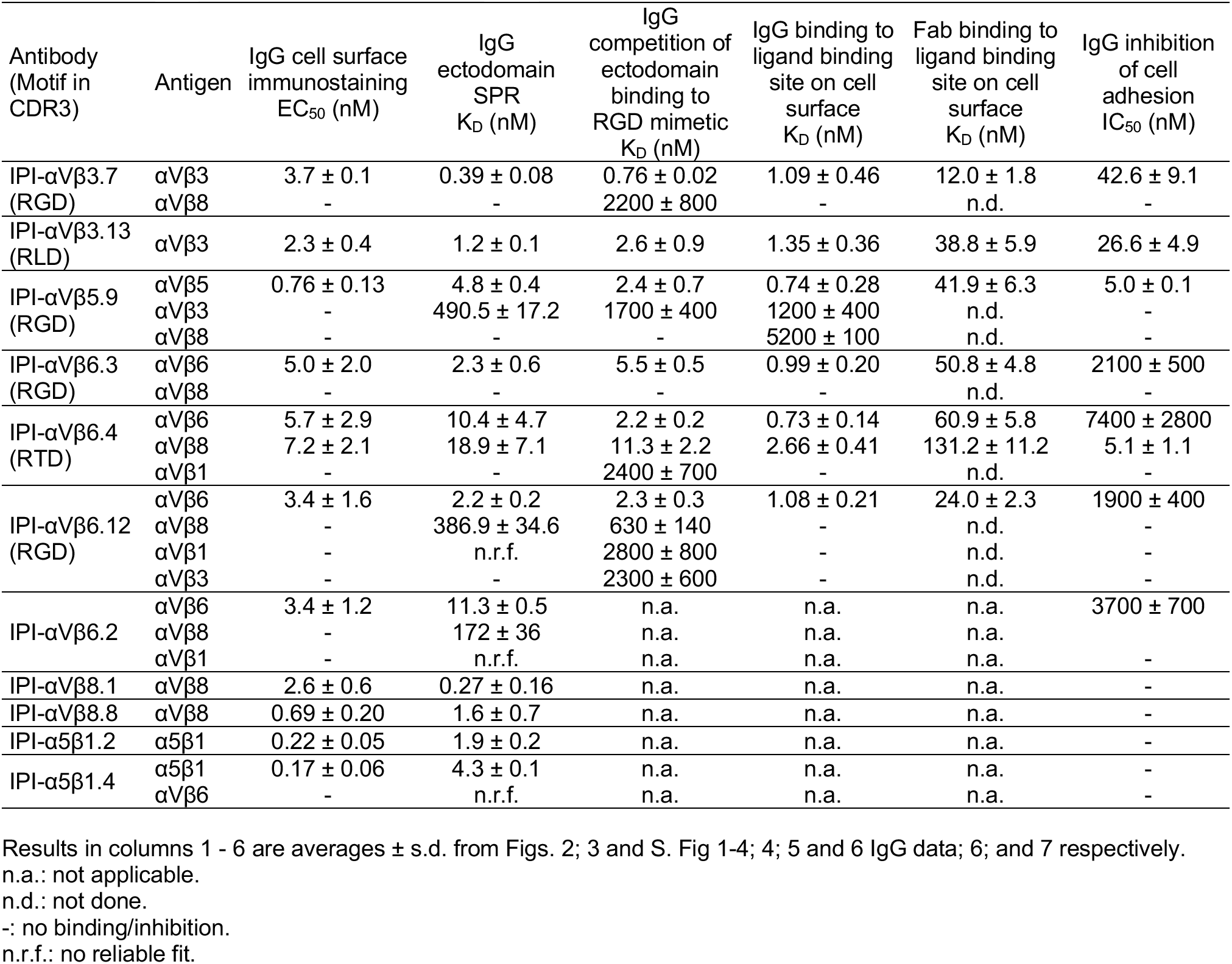
Binding characteristics of IPI anti-integrin antibodies.

The majority of the antibodies (6 out of the 11) block binding of small RGD mimetic ligands to their targeted integrin. Of the hundreds of previously described anti-integrin antibodies obtained by species-specific immunization, we know only a few with this characteristic: PAC-1, which has an RGD motif^28^ and mAb16^18,12^, which also has an RGD motif (unpublished). Despite the large number of integrins that recognize RGD motifs, we have selected for antibodies that are remarkably integrin-specific. Thus, IPI-αVβ3.13 was completely specific for integrin αVβ3, both in human and mouse. A previously described antibody, LM609, is specific for αVβ3 in human but as a mouse antibody does not react with mouse αVβ3^23^ and also does not block binding of small RGD mimetics to integrins (unpublished). IPI-αVβ6.4, with an RTD sequence, crossreacts between αVβ6 and αVβ8 with similar affinity. The other four antibodies, all with RGD sequences bound with low nanomolar affinities to their target integrins and showed greater than 100-fold higher affinity for the target than for any other integrin. Specificity of the six antibodies with RGD-like motifs is likely to be imparted by binding to regions outside of the RGD-binding pocket as well as by the presence of an RTD or RLD sequence in place of RGD in two of them. These antibodies will have many applications in the integrin field as ligand-binding blocking reagents, including the antibodies that show cross-reactivity, because we have defined their K_D_ and EC_50_ values (Table 2). Using these values, the percent bound equals 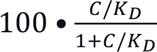

As an example, we know of no previously defined inhibitory αVβ5 antibody. By using IPI-αVβ5.9 IgG at 8.7 nM (10x its K_D_ for competing binding of a ligand to cell surface αVβ5), it would inhibit 90% of ligand binding to αVβ5 while inhibiting <1% of binding to cell surface αVβ3 or αVβ8. Furthermore, by using it at 50 nM (10x its IC_50_ for inhibiting cell adhesion), it essentially completely blocks all adhesion.

The EC_50_, IC_50_, and K_D_ values in Table 2 show several trends. By competing RGD mimetic binding, affinities of IgG for the ectodomain are higher than affinities of Fab for the intact integrin on cell surface. Both measure monomeric interactions. Measurements using biological ligands for integrin α5β1 and α4β1 show the same trend; ensemble affinities are lower for cell surface integrins because their content of the high affinity extended-open conformation is lower than for ectodomain preparations ^26;29^. On the other hand, the IgG affinity for ectodomain determined with SPR and competitive binding with RGD-mimetic agree well with one another. This agreement demonstrates the reliability of our reported affinities. Yet another comparison, of IgG and Fab binding to the integrin ligand-binding site on the cell surface, shows the difference between divalent and monomeric binding. Direct comparisons in Fig. 6 show a 20 to 60-fold increase in effective affinity for IgG. A caveat is that these measurements are on overexpressing transfectants, and IgG affinity will be lower at lower integrin expression levels. Limited experience of staining tumor cell lines shows that immunostaining EC_50_ values are cell line-dependent (Fig. S8). Mn^2+^ can substantially increase integrin affinity for ligand and can enhance immunostaining of the RGD mimetic antibodies (Fig. S8E). Among the assays for crossreactivity, competition assays were the most sensitive because a single concentration of the FITC-labeled RGD mimetic is used and the competitor can cover a broader range of concentrations. In contrast, in immunostaining and SPR, the background signal increases with the concentration of the antibody or antigen, respectively.

The αV subunit is unique among integrin α subunits in associating with five different β subunits, three of which, β5, β6, and β8, associate only with αV. Pair-wise competition between β-subunits revealed the order of preference to be αVβ6= αVβ8> αVβ3> αVβ1= αVβ5. A limitation is that although we used native β-subunit cDNAs all expressed in the same vector, we assumed β-subunit precursor expression was identical. However, we verified the same trend in several native tumor cell lines. Earlier, we found that the BJ-5a fibroblast cell line expresses integrins α5β1, αVβ1, αVβ3, and αVβ5^12^. Further cell lines studied here show that even when αV and β1 subunits are abundant, αVβ1 is not expressed when the more dominant αVβ3 and αVβ8 (LN229) or αVβ6 integrins (HT29) are expressed. However, both cell types expressed αVβ5, which appears to compete similarly to αVβ1 for the αV subunit in transfectants.

Expression of αVβ5 but not αVβ1 by these cells suggests that the αV subunit of αVβ1 also competes poorly for the β1 subunit with the other 11 α-subunits that associate with β1. In zebrafish integrins, a trend similar to that seen here was found in which αV associated less well with the β1-subunit than with the β3, β5, and β6-subunits^30^. During divergence among integrin orthologues in vertebrate evolution, both the αV and β1 subunits face the dilemma of retaining association with a larger number of β and α-subunits than any other integrin subunit. Nonetheless, our data suggests that the β1 subunit competes as effectively as β5 for αV in transfectants, despite the ability of the β1 and β5-subunits to associate with a total of 12 and 1 α-subunits, respectively.

## Methods

### Expression of full-length integrin on the cell surface

cDNA encoding native integrin α and β-subunits from Genscript (gene and accession No. are: hITGAV, NM_002210.5; hITGB1, NM_002211.3; hITGB3, NM_000212.3; hITGB5, NM_002213.5; hITGB6, NM_000888.5; hITGB8, NM_002214.3; hITGA2B, NM_000419.5; mITGA2B, NM_010575; mITGB8, NM_177290.3) and Sino Biological (gene and accession No. are: hITGA8, NM_003638.1; hITGA5, NM_002205.2; mITGAV, NM_008402.2; mITGA5, NM_010577.2; mITGB1, NM_010578.1; mITGB3, NM_016780.2; mITGB5, NM_010580.2; mITGB6, NM_021359.2; mITGA8, NM_001001309.2) were amplified by PCR and inserted into the pD2529 CAG vector (ATUM). The native signal sequence was replaced with an N-terminal CD33 secretion peptide (MPLLLLLPLLWAGALA), followed by full-length sequence. For mouse α-subunits only, the full-length sequence was followed by a P2A sequence and GFP. All β-subunit full-length constructs were followed by a P2A sequence (ATNFSLLKQAGDVEENPGP) and mCherry. The α and β cDNAs were transiently transfected into Expi293 α5^-^/αV^-^ cells ^31^ using FectoPro (Polyplus) according to the manufacturer’s instructions. After 24 hours of transfection, 3 mM valproic acid and 4g/L of glucose were added. Cells were used 48 hours after transfection.

### Expression and purification of integrin ectodomains

Ectodomains utilized the same full length sequences, truncated before the transmembrane domain. The α-subunit ectodomain sequence was followed by a HRV3C cleavage site (LEVLFQG), acid coil (AQCEKELQALEKENAQLEWELQALEKELAQ), Protein C tag (EDQVDPRLIDGK), and Strep twin tag (SAWSHPQFEKGGGSGGGGGSAWSHPQFEK). The β-subunit ectodomain was followed by HRV3C cleavage site, basic coil (AQCKKKLQALKKKNAQLKWKLQALKKKLAQ), HA tag (YPYDVPDYA), deca-histidine tag, P2A sequence, and mCherry. 7 days after transfection and supplementation as described above, supernatants were harvested and purified using His-Tag purification resin (Roche, cOmpelte^TM^, Cat No.5893682001), followed by size-exclusion chromatography in 20 M Hepes or Tris pH 8, 150 mM NaCl, 1 mM CaCl_2_, and 1 mM MgCl_2_ (GE Healthcare, AKTA purifier, Superdex 200). The clasped integrin ectodomains were concentrated to ∼1 mg/mL, flash frozen in liquid nitrogen and stored at -80°C.

### K562 stable transfectants expressing full-length RGD-binding integrins

For αVβ1, αVβ3, αVβ5, αVβ6, and αVβ8, αIIbβ3, and α8β1 transfectants, the appropriate full-length plasmids described above were electroporated into K562 cells, which express α5β1 as the sole RGD-binding integrin. Transfectants were selected with 3 µg/mL puromycin. αV transfectants were further FACS sorted using Alexa488-17E6 (anti-αV) and mCherry. α8β1 and αIIbβ3 transfectants were further FACS sorted using mCherry.

### Kinetic measurements using SPR

High-throughput SPR binding kinetics experiments used a Carterra LSA instrument with an HC-30M chip (Carterra-bio, catalog#4279) with a 384-ligand array format. The experiment was setup according to Carterra’s standard protocol. Briefly, antibodies were captured using immobilized goat anti-human IgG Fc secondary antibody (Jackson Immuno laboratory, catalog#109-005-098). A two-fold dilution series ranging from 0.07825 nM to 12.5 nM of purified integrin ectodomains as analyte in 10 mM HEPES (pH 7.5), 150 mM NaCl, 1 mM MgCl_2_, 1 mM CaCl_2_, 0.05% Tween 20, and 0.5 mg/mL BSA was sequentially injected (capture kinetics). After each 5 min association phase and 5 min dissociation phase, the association phase for the next highest concentration began.

Instrument software was used to subtract the reference cell background and for Y-alignment. Data were then globally fitted with two equations in Prism with shared *k_on_, k_off_,* and R_max_:

For the association phase, when *t* (time) is smaller than *t*_d_ (dissociation start time):

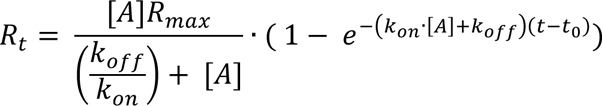

For the dissociation phase, when *t* (time) is larger than *t*_d_ (dissociation start time):

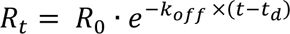

where *R_t_* is the observed response at time t, [*A*] is the analyte concentration, *k_off_* is the off-rate and *k_on_* is the on-rate, R_max_ is the maximal SPR response. R_max_ is a fitting parameter defined using the targeted integrin analyte for each integrin antibody and is used globally with all other integrin analytes binding to that antibody. *t_0_* is the fitted start time of each cycle and is used to calculate the initial response units at the beginning of each new association phase. *R_0_* is *R_t_* at *t*=*t_d_*.

Prism input is as follows:

ligand=HotNM*1e-9
Kob=[ligand]*Kon+Koff
Kd=Koff/Kon
Eq=Bmax*ligand/(ligand + Kd)
Association=Eq*(1-exp(-1*Kob*(X-t0)))
YatTime0 = Eq*(1-exp(-1*Kob*Time0))
Dissociation= YatTime0*exp(-1*Koff*(X-t0-Time0))
Y=IF(X<Time0, Association, Dissociation) + NS
X: Time
Y: Total binding
Koff: Dissociation constant in inverse time units.
Kon: Association constant in inverse time multiplied by inverse concentration.
Kd: Computed from Koff/Kon, in Molar units.
t0 is used to correct for the experiment start time and to compensate for the initial response units due to the previous binding cycle.
Bmax: Maximum binding at equilibrium with maximum concentration of analyte, in units of Y axis.
HotNM (the concentration of analyte in nM)
Time0 (the time at which dissociation was initiated).
NS = 0.

### Antibodies and fluorescent labeling

Antibodies were 17E6 (anti-αV)^16^, mab16 (anti-α5)^18^, mab13 (anti-β1)^18^, LM609 (anti-αVβ3)^23^, 7.1G10 (anti-αVβ6)^32^, ADWA11 (anti-αVβ8)^33^, and Biogen-αVβ1.Ab5 (anti-αVβ1) SEQ ID NO:35^25^.

Alexa Fluor 647 NHS Ester (Thermo Fisher Scientific, A20006) was used to directly label integrin antibodies following the manufacturer’s protocol. Briefly, 1 mg of antibody (5 mg/mL) was incubated with 10 μg of Alexa Fluor 647 NHS Ester (10 μg/μL in DMSO) in PBS pH 7.4 at room temperature for 1 hr in the dark. Unconjugated dye was removed by size exclusion chromatography (GE Healthcare, AKTA purifier, Superdex 200). IgG concentration was calculated as:

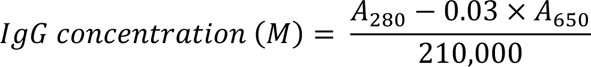

The dye ratio was calculated as

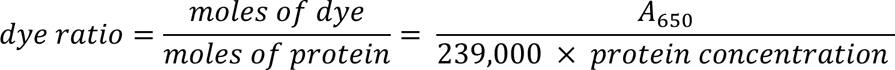

### Indirect immunofluorescent flow cytometry

K562 stable transfectants expressing human RGD-binding integrins or K562 WT cells endogenously expressing α5β1 or Expi293F α5^-^/αV^-^ mouse integrin transfectants (10^6^ cells/mL) were incubated with the indicated concentration of antibodies in Hanks’ balanced salt solution (HBSS) with 20 mM HEPES pH 7.4, 1% BSA, 1 mM Ca^2+^, and 1mM Mg^2+^ (or 1mM Mn^2+^ when indicated) for 1hr on ice followed by three washes. Cells were then stained with APC-conjugated goat anti-human IgG (Jackson Immuno Research, Catalog 109-135-098) at a 1:150 dilution, followed by three washes, and subjected to FACS (BD FACSCanto II). The background mean fluorescence intensity (MFI) was determined using a human IgG1 isotype control (Bioxcell #BE0297) at the same concentration as the primary antibodies. Data analysis used FlowJo (Version 10.7.1).

LN229 (ATCC CRL-2611) and HT29 (ATCC HTB-38) cells were stained identically with first antibodies at 50 nM, except for rat and mouse antibodies, Alexa Fluor 647 goat anti-rat IgG (Invitrogen, catalog A-21247) at 2 μg/mL and Alexa Fluor 647 goat anti-mouse F(ab’)2 (Invitrogen, catalog A-21237) at 2 μg/mL were used, respectively. Background mean fluorescence intensity (MFI) was determined using rat IgG2a, BD Catalog 553933 and mouse IgG, clone X63; human IgG Bioxcell #BE0297.

### Fluorescence polarization

FITC-labeled aminocaproic acid-disulfide-cyclized ACRGDGWCG peptide (FITC-cyclic-ACRGDGWCG) and FITC-labeled aminocaproic acid-GRGDLGRLKK peptide (FITC-proTGFβ3 peptide) were synthesized by GenScript. Preliminary experiments (Supplementary Fig. 5) were with 10 nM of FITC-labeled peptide probe and indicated integrin ectodomain concentrations in 10 mM HEPES pH 7.5, 150 mM NaCl, 1 mM MgCl_2_, 1 mM CaCl_2_, and 0.5 mg/mL BSA (10 µL). The mixture was allowed to equilibrate for 2 hr in the dark and the FP signal was measured by Synergy NEO HTS multi-mode microplate reader (Biotek). The background FP signal was measured by supplementing the reaction with 10 mM EDTA. Affinities were obtained by fitting the curve to previously published equations^26^ (Supplementary Equation S17).

For the competition assays, samples (10 µL) contained 10nM FITC-cyclic-ACRGDGWCG or FITC-proTGFβ3 peptide, integrin ectodomain, and antibodies at indicated concentrations in the same buffer and condition as described above. Data were fitted globally using previously developed equations^26^ (Supplementary Equation S28), with the maximum FP value in the absence of antibody and the minimum FP value as shared parameters, and affinities for each titrator as individual parameters. The α5β1 minibinder used in this experiment was obtained using a method similar to that described in ^9^.

### IgG and Fab binding to integrin ligand-binding sites on the cell surface

The affinity of antibodies to integrins αVβ1 and α8β1 expressed on K562 stable transfectants, as well as α5β1 expressed on K562 wild-type cells, was measured by enhancement of binding of 10nM AF647-9EG7 Fab. Cells (10^5^ in 100 µL) were mixed with 10 nM AF647-9EG7 Fab and indicated concentrations of antibodies or cyclic-ACRGDGWCG in L15 medium containing 1% BSA for 2 hrs at room temperature. Flow cytometry was without washing to ensure that values were obtained under equilibrium conditions. The MFI values of AF647-9EG7 Fab in the presence of various concentrations of titrators on each cell line were fitted by a three-parameter dose-response curve. The errors for the affinities are the difference from the mean value from duplicate experiments.

To determine the affinity of FITC-proTGFβ3 peptide to αVβ6 and αVβ8 on the K562 cell surface (Figure S6), 100 µL of cells (10^6^/mL) were mixed with indicated concentrations of FITC-proTGFβ3 peptide in L15 medium containing 1% BSA for 2 hrs at room temperature and subjected to flow cytometry without washing. Background fluorescence was measured with 10 mM EDTA in the binding buffer. The background-subtracted mean fluorescence intensity (MFI) at each concentration of FITC-proTGFβ3 peptide was fitted to a three-parameter dose-response curve for K_d_, background MFI, and maximum MFI.

The affinities of cRGDfk peptide with lysine side chain conjugated to TideFluor5WS (TF5WS-cRGDfk) to αVβ3 (K_d_ = 57 ± 6 nM) and αVβ5 (K_d_ = 51 ± 8 nM) on cell surface were previously determined^12^. The binding affinity of FITC labeled Echistatin (FITC-Echistatin) to αIIbβ3 (K_d_ = 248 ± 14 nM) was previously quantified^31^.

IgG and Fab affinities for intact αVβ3, αVβ5, αVβ6, αVβ8, and αIIbβ3 on K562 stable transfectants were measured by competing fluorescently labeled RGD-containing peptidomimetics. Cells (10^6^/mL in 100 µL) were mixed with the indicated probe and antibody concentration in L15 medium with 1% BSA. After 2 hrs in the dark at room temperature to ensure equilibrium, cells were subjected to FACS.

### Cell adhesion assays

50 µL of ligands in PBS (pH 7.4) were coated to ELISA high binding 96-well plates (Corning, REF 3590) at 4°C for 16 hrs. Plates were washed and blocked for 1hr at 37°C with PBS containing 3% BSA. Integrin transfectants in L15 medium (10^6^ cells/mL in 50 µL) were mixed with antibodies in 50 µL in L15 medium and added to wells. After 1 hr at 37°C, the fluorescent intensity of mCherry, which was co-expressed with the transfected β-subunit through self-cleaving P2A peptide (Methods, section 2), was detected at 625 nm using Biotek Synergy NEO HTS multi-mode microplate reader. After three washes by gently removing the L15 medium and replenishment with 100 µL of L15 medium, the plate was read again to obtain the fraction of cells bound. For αVβ6 and αVβ8 transfectants, cells and antibodies were pre-incubated for 1 hr and 37°C, before adding to wells.

### Competition between integrin β-subunits for the αV-subunit

Integrin α and β-subunits were transfected as described above for cell surface expression using 1.8 μg plasmid per 1.8 mL of cells (3×10^6^/mL). The experiments are described in detail in Supplementary Fig. 7 and Fig. 8.

Expi293F αV^-^/α5^-^ transfectants (5 x 10^4^ in 50 µL) were stained with directly Alexa 647-labeled integrin antibodies at 100 nM or Alex647-labeled 17E6 anti-αV at 40 nM in Hanks’ balanced salt solution, 20mM HEPES, 1mM Ca^2+^, 1mM Mg^2+^ and 1% BSA on ice for 1 hr and subjected to FACS after 3 washes.

Background was measured using Alexa 647-labeled human natalizumab (anti-α4) for human antibodies or Alexa 647-labeled mouse IgG1 (clone X63 isotype control) for 17E6 anti-αV and P1F6 (anti-β5). The specific MFI reported in Fig. S7 was background corrected as:

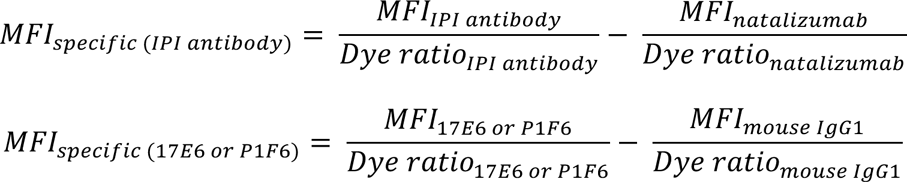

Due to variations in kinetics among different antibodies, the specific MFI cannot be directly compared between each integrin β-subunit antibody. To enable a direct comparison, a coefficient was calculated to adjust each β-subunit antibody MFI value relative to the MFI value of the 17E6 αV antibody using the equation:

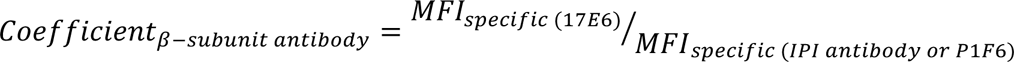

The calculated coefficient for each β-subunit antibody is indicated on each panel in Fig. S7. The MFI for each integrin shown in Fig. 8 is calculated as:

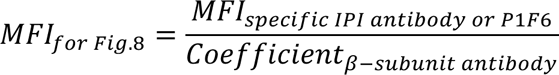

## Acknowledgements

This work was supported by NIH grants HL159714 and HL131729. We thank Xinru Wang from David Baker’s lab for providing α5β1 mini binder.

## Author contribution

Yuxin Hao, Conceptualization, Formal analysis, Investigation, Methodology, Validation, Writing – original draft, Writing – review and editing; Jiabin Yan, Formal analysis, Investigation; Courtney Fraser, Formal analysis, Investigation; Aiping Jiang, Formal analysis, Investigation; Murali Anuganti, Investigation, Methodology; Roushu Zhang, Investigation; Joseph Jardine, Methodology, Investigation; Rob Meijers, Supervision; Jing Li, Conceptualization, Supervision, Writing – review and editing; Timothy A. Springer, Conceptualization, Funding acquisition, Project administration, Supervision, Writing – review and editing.

## Supplementary figure legends

**Supplementary figure 1.**
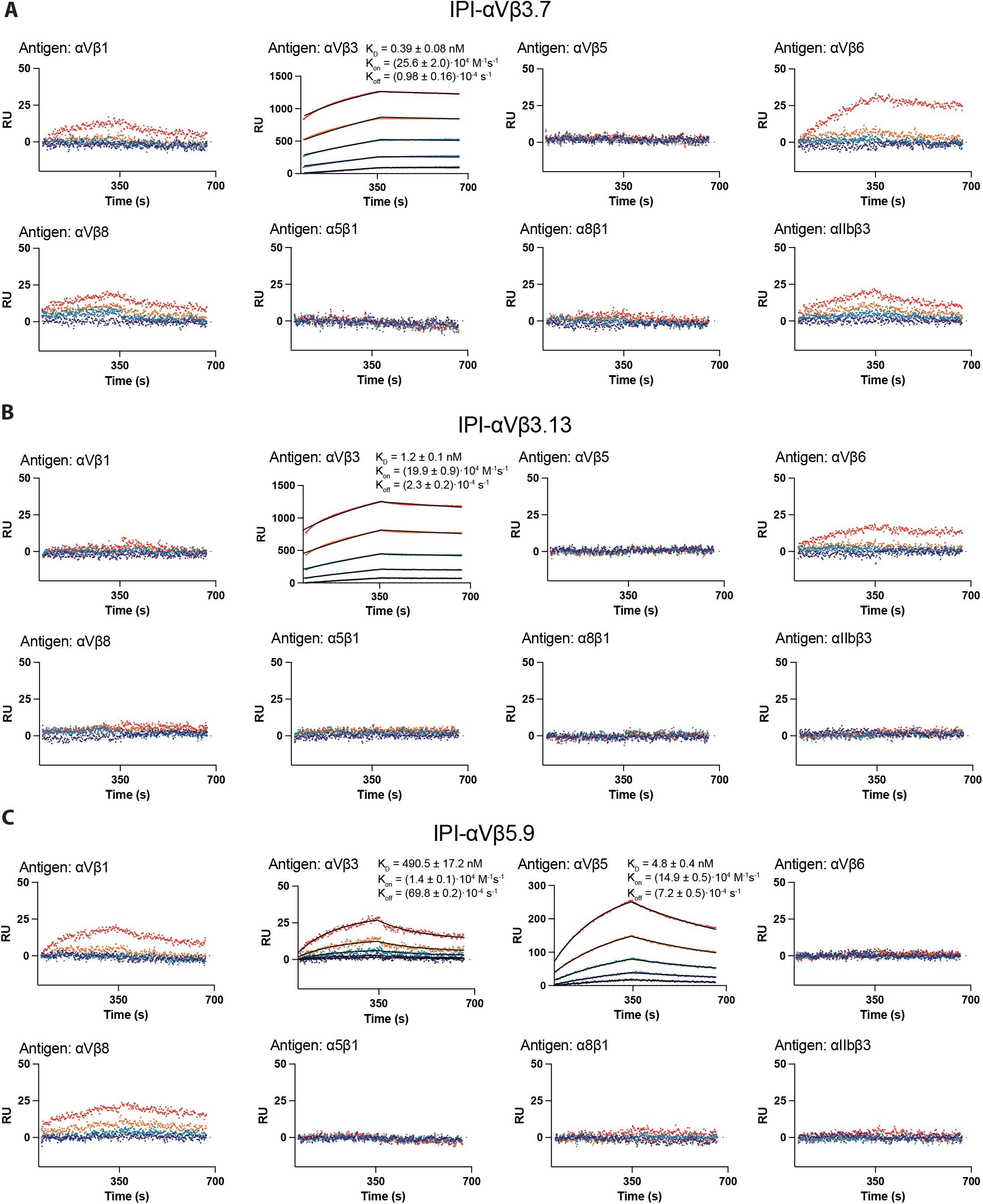
SPR sensorgram of binding of the soluble ectodomain (0.78, 1.56, 3.12, 6.25, and 12.50 nM) of each RGD-binding integrin to each immobilized IPI antibody (in Supplementary Figures 1-4). Fits (thin black lines) are shown when valid. Related to main Figure 3.

**Supplementary figure 2.**
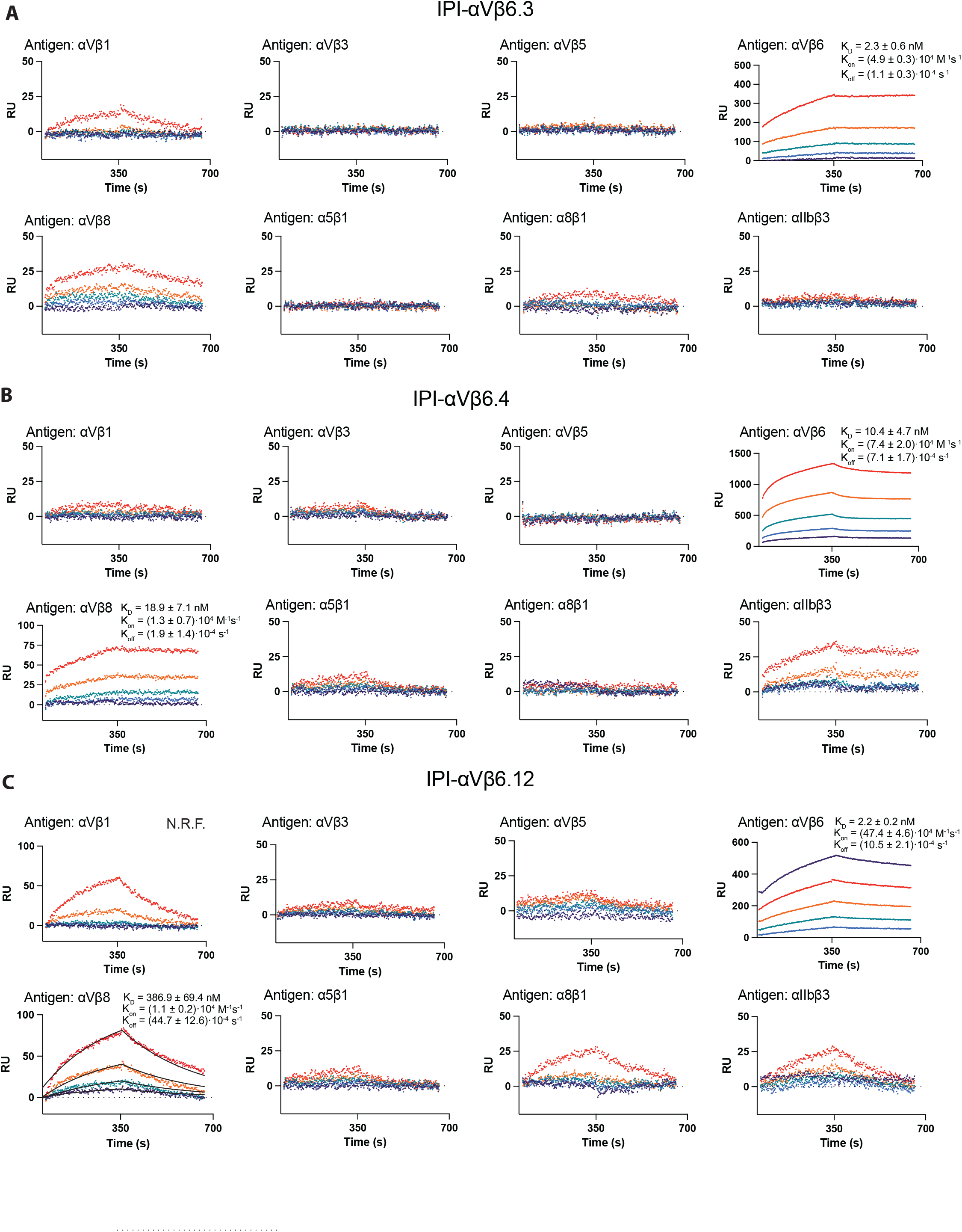
See Supplementary figure 1 legend.

**Supplementary figure 3.**
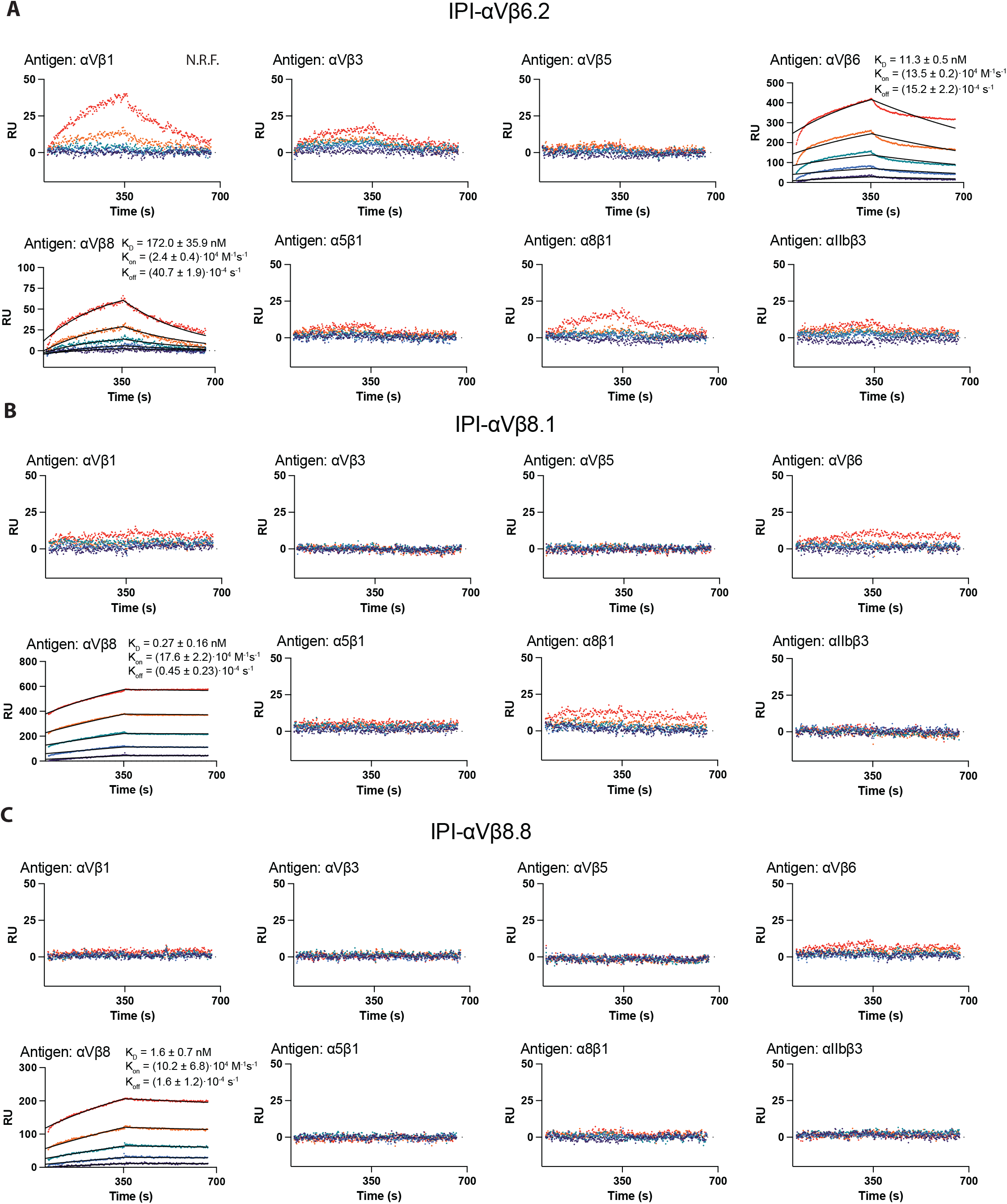
See Supplementary figure 1 legend.

**Supplementary figure 4.**
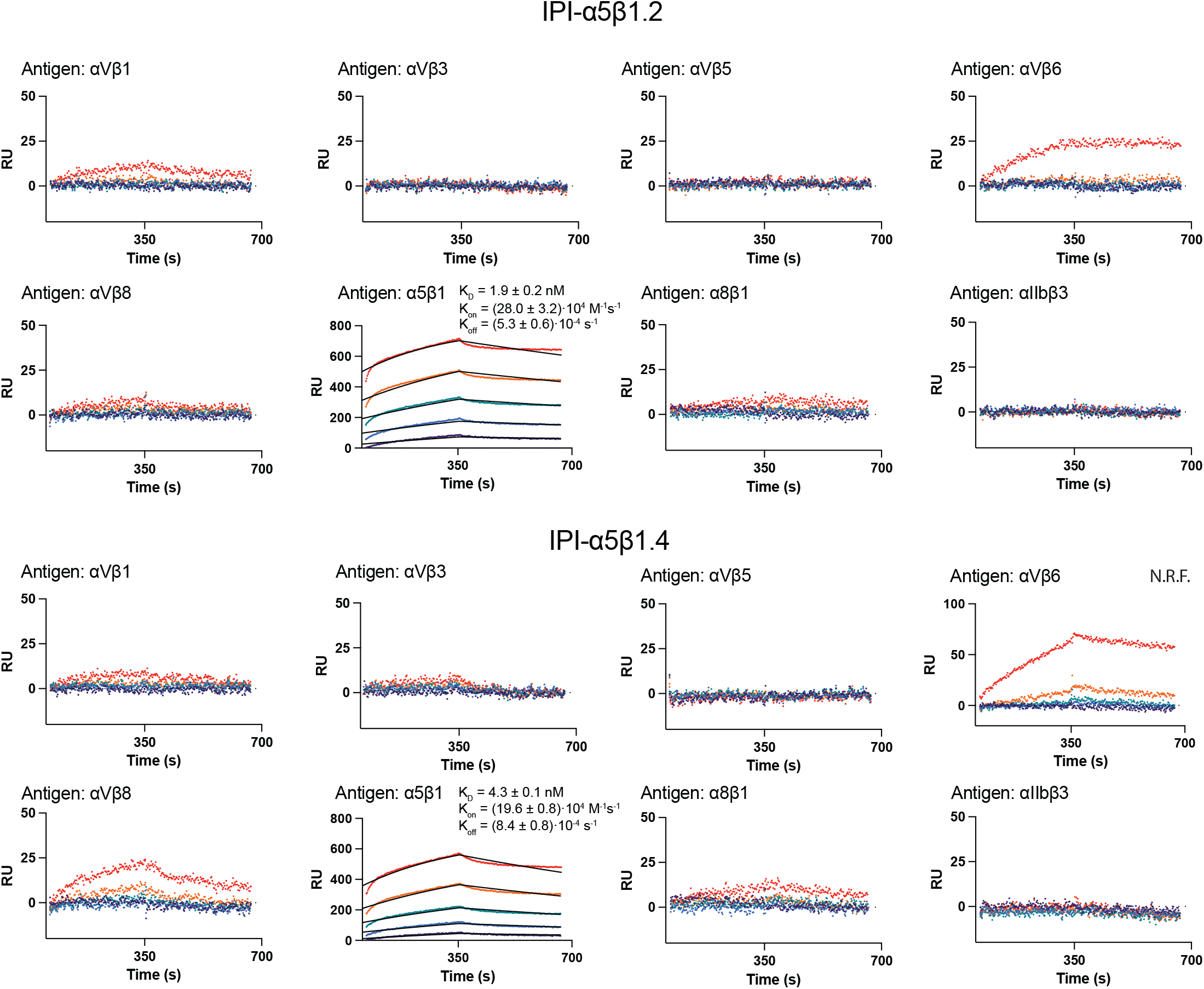
See Supplementary figure 1 legend.

**Supplementary figure 5.**
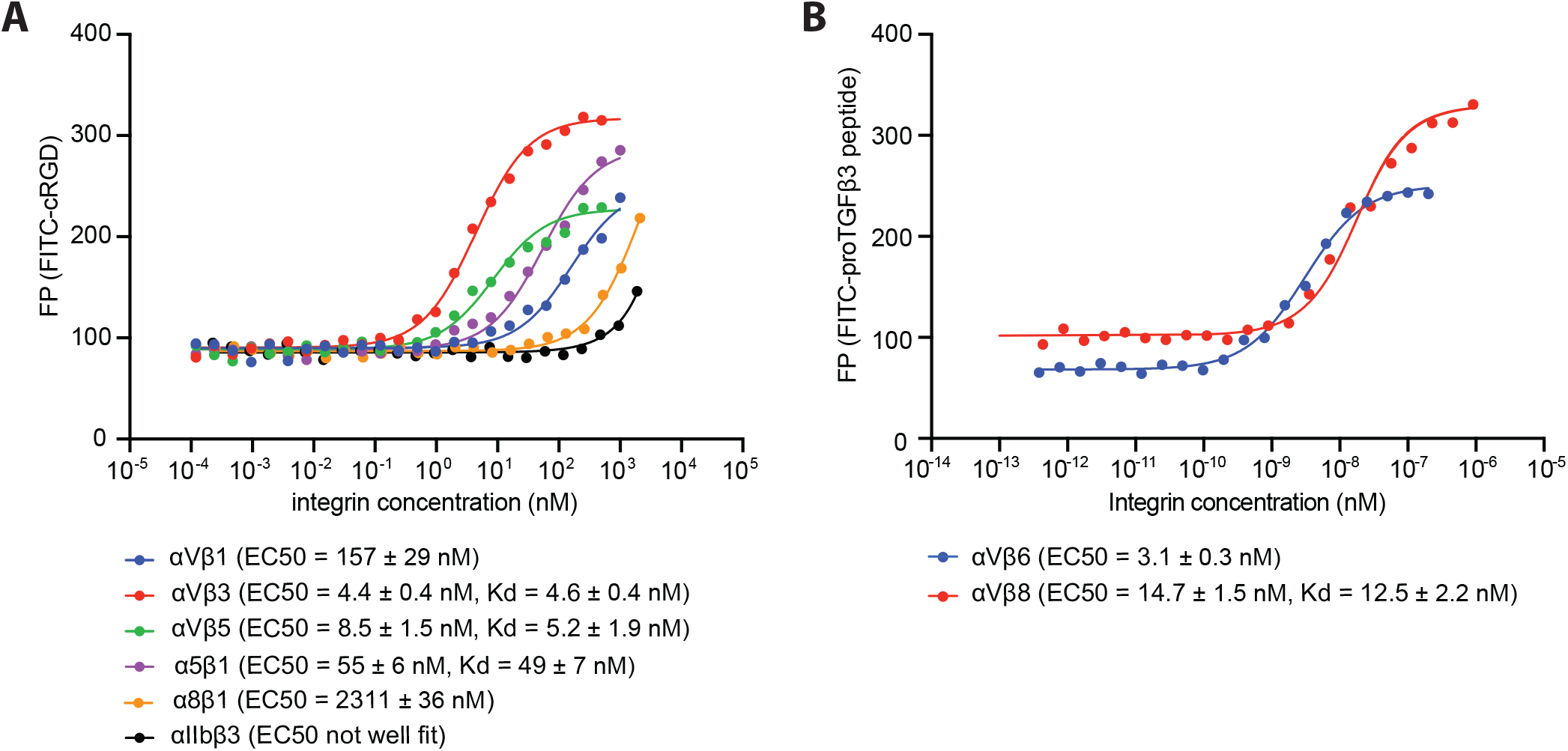
Titrations of integrin ectodomain binding to FITC-labeled peptidomimetics in fluorescence polarization (used to determine the integrin concentrations used in main Figure 4). (A) Binding of integrin ectodomains to FITC-cyclic-ACRGDGWCG. (B) Binding of αVβ6 and αVβ8 ectodomains to FITC-proTGFβ3 peptide. Binding was in 10 mM HEPES pH 7.5, 150 mM NaCl, 1 mM MgCl_2_, 1 mM CaCl_2_, and 0.5 mg/mL BSA. Background was measured in binding buffer supplemented with 10 mM EDTA. Background-subtracted FP was fitted to a three-parameter dose-response curve to obtain EC50. Means and errors are from the nonlinear least square fits. When good fits were obtained, K_D_ values are reported from fitting the saturation binding equations published previously (Methods).

**Supplementary Figure 6.**
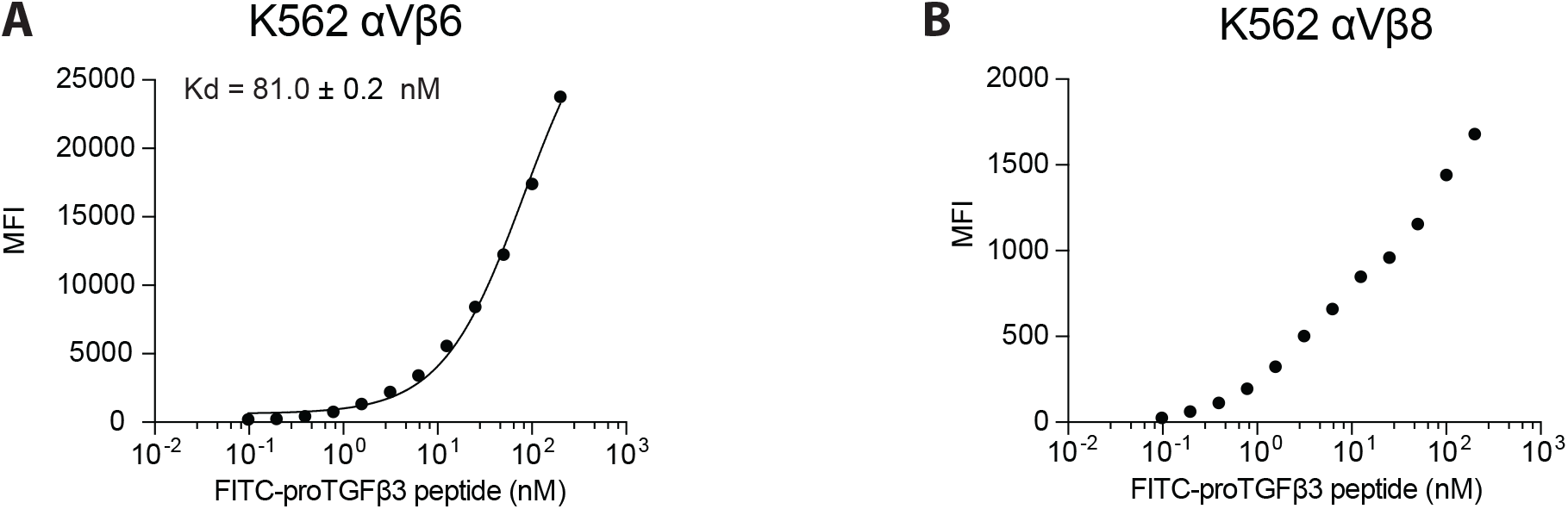
Binding of FITC-proTGFβ3 peptide for intact αVβ6 (A) and αVβ8 (B) on cell surfaces (used to determine FITC-proTGFβ3 peptide concentrations used in main Figure 5 and 6). Binding of FITC-proTGFβ3 peptide was in L15 medium containing 1% BSA and used flow cytometry without washing. Background was measured in binding buffer supplemented with 10 mM EDTA. Background-subtracted MFI at each FITC-proTGFβ3 peptide concentration was fitted to three three-parameter dose-response curve. The errors for the EC50 values are the difference from the mean of duplicate experiments. The K_D_ of FITC-proTGFβ3 peptide to αVβ8 was hard to quantify due to its low affinity, resulting in a low signal-to-noise ratio when used at high concentrations of fluorescence-labeled peptide.

**Supplementary Figure 7.**
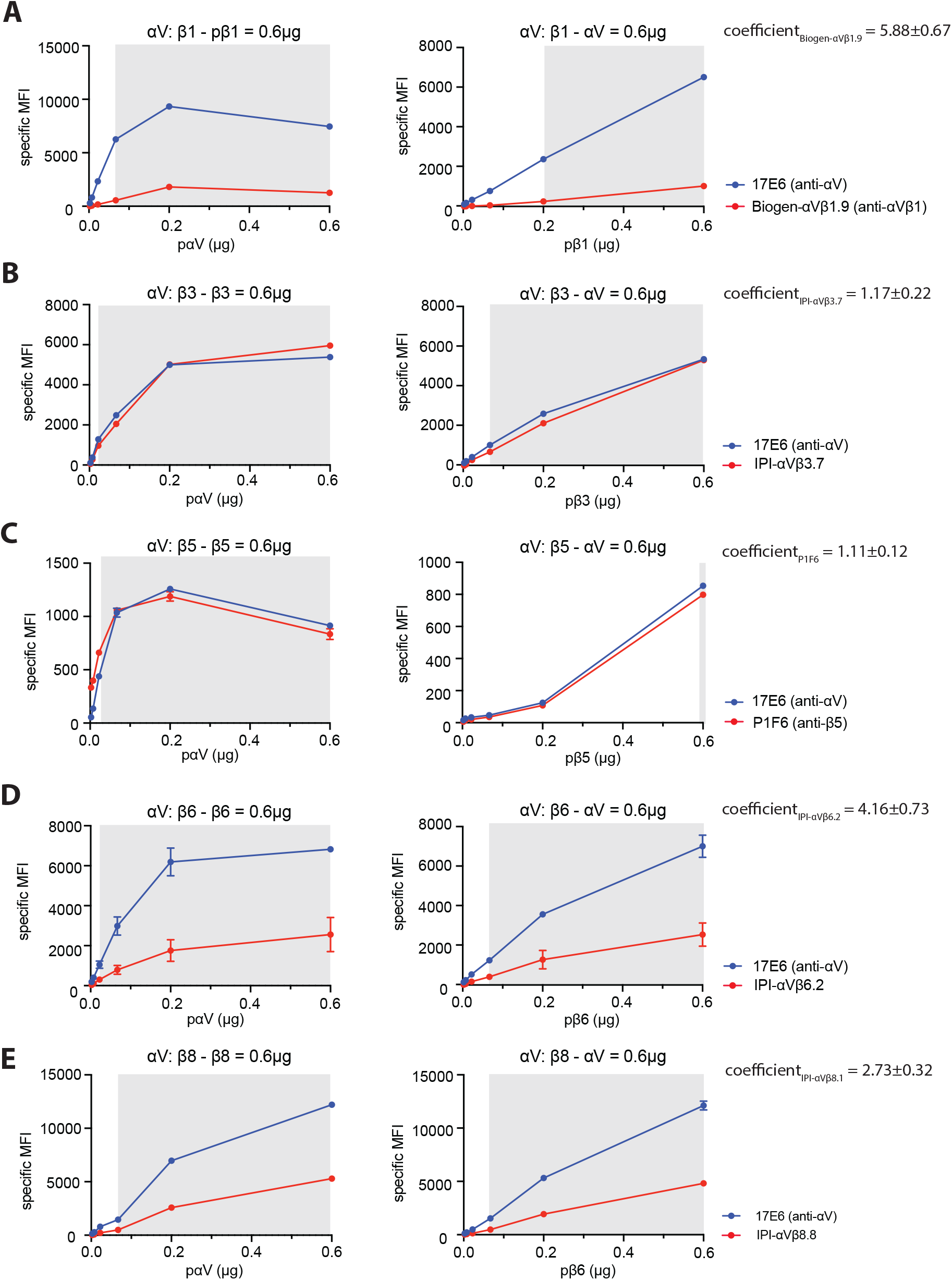
Titration between integrin αV-subunit and β-subunits. In each titration, the concentration of the αV-subunit plasmid (pαV) or β-subunit plasmid (pβ) remained constant at 0.6 µg, while β-subunit plasmid or αV-subunit plasmid, respectively, was titrated until reaching 0.6 µg. In all reactions, empty vector plasmid was added to make the total plasmid concentration 1.2 µg. MFI of directly fluorophore-labeled integrin antibodies was measured by flow cytometry and was normalized by the dye labeling ratio of each antibody. The coefficient of each β-subunit antibody used to normalize MFI relative to the MFI of the 17E6 αV antibody in Fig. 8 is indicated on the upper right of each panel (using data points included in the gray area, as low MFI data can be influenced by endogenous β subunits in the cells). The reported value is the mean and standard deviation from the data points in the gray area.

**Supplementary Figure 8.**
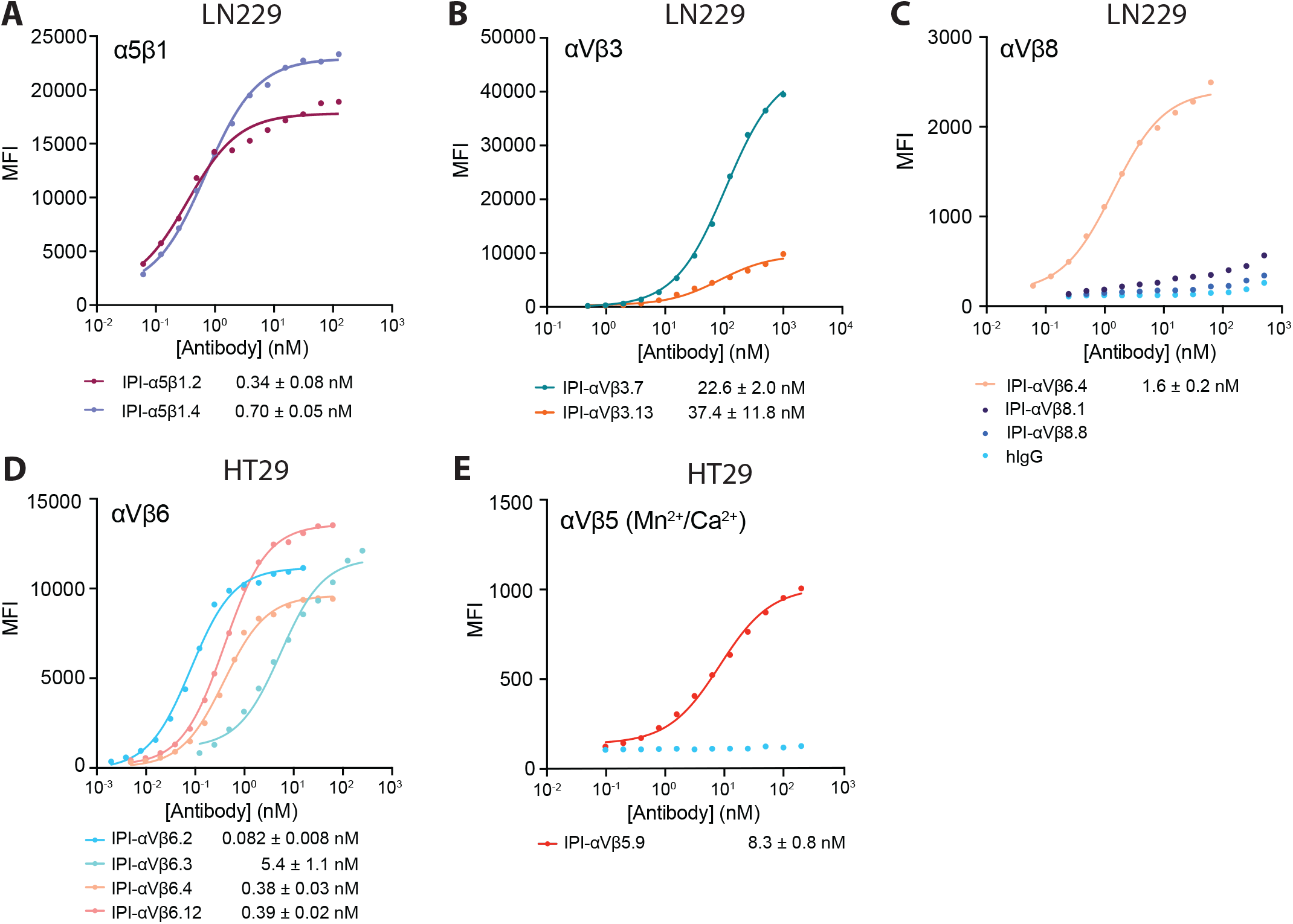
Indirect immunofluorescent staining of cell surface-expressed integrins on LN229 cells (A-C) and HT29 cells (D, E). Cells were stained with indicated concentrations of integrin antibodies in HBSS buffer containing 1 mM Ca^2+^ and 1 mM Mg^2+^ except for IPI-αVβ5.9, which used 1 mM Mn^2+^ and 0.2 mM Ca^2+^. After washing, integrin antibodies were detected using APC-conjugated goat anti-human secondary antibodies and flow cytometry. The MFI at each antibody concentration after subtraction of isotype control at the same concentration was fitted to a three-parameter dose-response curve for EC_50_, background MFI, and maximum MFI; curves are only shown for antibodies with meaningful staining. The errors for the EC_50_ values are the standard errors from the non-linear least square fits.

**Supplemental Table 1.** KD and kinetic rates of IPI integrin antibodies.

| **Supplemental Table 1. $K_D$ and kinetic rates of IPI integrin antibodies.** |  |  |  |  |
| --- | --- | --- | --- | --- |
| | | $k_{on}$ ($10^4$ M<sup>-1</sup>s<sup>-1</sup>) | $k_{off}$ ($10^{-4}$ s<sup>-1</sup>) | $K_d$ (nM) |
| $\alpha V\beta 1$<br>ectodomain | IPI-$\alpha V\beta 3.7$ | - | - | - |
| | IPI-$\alpha V\beta 3.13$ | - | - | - |
| | IPI-$\alpha V\beta 5.9$ | - | - | - |
| | IPI-$\alpha V\beta 6.2$ | N.R.F. | N.R.F. | N.R.F. |
| | IPI-$\alpha V\beta 6.3$ | - | - | - |
| | IPI-$\alpha V\beta 6.4$ | - | - | - |
| | IPI-$\alpha V\beta 6.12$ | N.R.F. | N.R.F. | N.R.F. |
| | IPI-$\alpha V\beta 8.1$ | - | - | - |
| | IPI-$\alpha V\beta 8.8$ | - | - | - |
| | IPI-$\alpha 5\beta 1.2$ | - | - | - |
| | IPI-$\alpha 5\beta 1.4$ | - | - | - |
| $\alpha V\beta 3$<br>ectodomain | IPI-$\alpha V\beta 3.7$ | $25.6 \pm 2.0$ | $0.98 \pm 0.16$ | $0.39 \pm 0.08$ |
| | IPI-$\alpha V\beta 3.13$ | $19.9 \pm 0.9$ | $2.3 \pm 0.2$ | $1.2 \pm 0.1$ |
| | IPI-$\alpha V\beta 5.9$ | $1.4 \pm 0.1$ | $69.8 \pm 2.1$ | $490.5 \pm 17.2$ |
| | IPI-$\alpha V\beta 6.2$ | - | - | - |
| | IPI-$\alpha V\beta 6.3$ | - | - | - |
| | IPI-$\alpha V\beta 6.4$ | - | - | - |
| | IPI-$\alpha V\beta 6.12$ | - | - | - |
| | IPI-$\alpha V\beta 8.1$ | - | - | - |
| | IPI-$\alpha V\beta 8.8$ | - | - | - |
| | IPI-$\alpha 5\beta 1.2$ | - | - | - |
| | IPI-$\alpha 5\beta 1.4$ | - | - | - |
| $\alpha V\beta 5$<br>ectodomain | IPI-$\alpha V\beta 3.7$ | - | - | - |
| | IPI-$\alpha V\beta 3.13$ | - | - | - |
| | IPI-$\alpha V\beta 5.9$ | $14.9 \pm 0.5$ | $7.2 \pm 0.5$ | $4.8 \pm 0.4$ |
| | IPI-$\alpha V\beta 6.2$ | - | - | - |
| | IPI-$\alpha V\beta 6.3$ | - | - | - |
| | IPI-$\alpha V\beta 6.4$ | - | - | - |
| | IPI-$\alpha V\beta 6.12$ | - | - | - |
| | IPI-$\alpha V\beta 8.1$ | - | - | - |
| | IPI-$\alpha V\beta 8.8$ | - | - | - |
| | IPI-$\alpha 5\beta 1.2$ | - | - | - |
| | IPI-$\alpha 5\beta 1.4$ | - | - | - |
| $\alpha V\beta 6$<br>ectodomain | IPI-$\alpha V\beta 3.7$ | - | - | - |
| | IPI-$\alpha V\beta 3.13$ | - | - | - |
| | IPI-$\alpha V\beta 5.9$ | - | - | - |
| | IPI-$\alpha V\beta 6.2$ | $13.5 \pm 0.2$ | $15.2 \pm 2.2$ | $11.3 \pm 0.5$ |
| | IPI-$\alpha V\beta 6.3$ | $4.9 \pm 0.3$ | $1.1 \pm 0.3$ | $2.3 \pm 0.6$ |
| | IPI-$\alpha V\beta 6.4$ | $7.4 \pm 2.0$ | $7.1 \pm 1.7$ | $10.4 \pm 4.7$ |
| | IPI-$\alpha V\beta 6.12$ | $47.4 \pm 4.6$ | $10.4 \pm 1.8$ | $2.2 \pm 0.2$ |
| | IPI-$\alpha V\beta 8.1$ | - | - | - |
| | IPI-$\alpha V\beta 8.8$ | - | - | - |
| | IPI-$\alpha 5\beta 1.2$ | - | - | - |
| | IPI-$\alpha 5\beta 1.4$ | N.R.F. | N.R.F. | N.R.F. |

| **Supplemental Table 1, *cont.* K<sub>D</sub> and kinetic rates of IPI integrin antibodies** |  |  |  |  |
| --- | --- | --- | --- | --- |
|  |  | k<sub>on</sub> (10<sup>4</sup> M<sup>-1</sup>s<sup>-1</sup>) | k<sub>off</sub> (10<sup>-4</sup> s<sup>-1</sup>) | K<sub>D</sub> (nM) |
| αVβ8<br>ectodomain | IPI-αVβ3.7 | - | - | - |
|  | IPI-αVβ3.13 | - | - | - |
|  | IPI-αVβ5.9 | - | - | - |
|  | IPI-αVβ6.2 | 2.4 ± 0.4 | 40.7 ± 1.9 | 172.0 ± 35.9 |
|  | IPI-αVβ6.3 | - | - | - |
|  | IPI-αVβ6.4 | 1.3 ± 0.7 | 1.9 ± 1.4 | 18.9 ± 7.1 |
|  | IPI-αVβ6.12 | 1.1 ± 0.2 | 44.7 ± 12.6 | 386.9 ± 34.6 |
|  | IPI-αVβ8.1 | 17.6 ± 2.2 | 0.45 ± 0.23 | 0.27 ± 0.16 |
|  | IPI-αVβ8.8 | 10.2 ± 6.8 | 1.6 ± 1.2 | 1.6 ± 0.7 |
|  | IPI-α5β1.2 | - | - | - |
|  | IPI-α5β1.4 | - | - | - |
| α5β1<br>ectodomain | IPI-αVβ3.7 | - | - | - |
|  | IPI-αVβ3.13 | - | - | - |
|  | IPI-αVβ5.9 | - | - | - |
|  | IPI-αVβ6.2 | - | - | - |
|  | IPI-αVβ6.3 | - | - | - |
|  | IPI-αVβ6.4 | - | - | - |
|  | IPI-αVβ6.12 | - | - | - |
|  | IPI-αVβ8.1 | - | - | - |
|  | IPI-αVβ8.8 | - | - | - |
|  | IPI-α5β1.2 | 28.0 ± 3.2 | 5.3 ± 0.6 | 1.9 ± 0.2 |
|  | IPI-α5β1.4 | 19.6 ± 0.8 | 8.4 ± 0.8 | 4.3 ± 0.1 |
| α8β1<br>ectodomain | IPI-αVβ3.7 | - | - | - |
|  | IPI-αVβ3.13 | - | - | - |
|  | IPI-αVβ5.9 | - | - | - |
|  | IPI-αVβ6.2 | - | - | - |
|  | IPI-αVβ6.3 | - | - | - |
|  | IPI-αVβ6.4 | - | - | - |
|  | IPI-αVβ6.12 | - | - | - |
|  | IPI-αVβ8.1 | - | - | - |
|  | IPI-αVβ8.8 | - | - | - |
|  | IPI-α5β1.2 | - | - | - |
|  | IPI-α5β1.4 | - | - | - |
| αIIbβ3<br>ectodomain | IPI-αVβ3.7 | - | - | - |
|  | IPI-αVβ3.13 | - | - | - |
|  | IPI-αVβ5.9 | - | - | - |
|  | IPI-αVβ6.2 | - | - | - |
|  | IPI-αVβ6.3 | - | - | - |
|  | IPI-αVβ6.4 | - | - | - |
|  | IPI-αVβ6.12 | - | - | - |
|  | IPI-αVβ8.1 | - | - | - |
|  | IPI-αVβ8.8 | - | - | - |
|  | IPI-α5β1.2 | - | - | - |
|  | IPI-α5β1.4 | - | - | - |

